# Back to the future: omnipresence of fetal influence on the human brain through the lifespan

**DOI:** 10.1101/2022.12.02.514196

**Authors:** Kristine B. Walhovd, Stine Kleppe Krogsrud, Inge K. Amlien, Øystein Sørensen, Yunpeng Wang, Anne Cecilie Sjøli Bråthen, Knut Overbye, Jonas Kransberg, Athanasia M. Mowinckel, Fredrik Magnussen, Martine Herud, Asta K. Håberg, Anders M. Fjell, Didac Vidal-Piñeiro

## Abstract

Human fetal development has been associated with brain health at later stages. It is unknown whether growth in utero, as indexed by birth weight (BW), relates consistently to lifespan brain characteristics and changes, and to what extent these influences are of a genetic or environmental nature. Here we show remarkably stable and life-long positive associations between BW and cortical surface area and volume across and within developmental, aging and lifespan longitudinal samples (N = 5794, 4-82 years of age, w/ 386 monozygotic twins, followed for up to 8.3 years w/12,088 brain MRIs). In contrast, no consistent effect of BW on brain changes was observed. Partly environmental effects were indicated by analysis of twin BW discordance. In conclusion, the influence of prenatal growth on cortical topography is stable and reliable through the lifespan. This early life factor appears to influence the brain by association of brain reserve, rather than brain maintenance. Thus, fetal influences appear omnipresent in the spacetime of the human brain throughout the human lifespan. Optimizing fetal growth may increase brain reserve for life, also in aging.

## Introduction

It is established that a substantial portion of functional variation through the lifespan, including in older age, is of neurodevelopmental origin^1–4^. Evidence converges on early life factors being important for normal individual differences in brain, mental health and cognition across the lifespan^5–9^, as well as risk of psychiatric^10^ and neurodegenerative disease in older age^11^. Obtaining reliable indicators of individual early life factors is a major challenge. In this regard, birth weight (BW) stands out as a solid available measure. BW reflects fetal and maternal genetic, but also other in utero environmental factors affecting fetal growth ^12–14^, including brain growth ^6,15–17^. By now, a series of studies have established that BW relates positively to mental health, cognitive function and brain characteristics, including neuroanatomical volumes and cortical surface area as measured in different age groups ^4–6,15,18,19^. However, it is unknown whether and how BW relates to brain characteristics through the lifespan, how consistent effects are within and across samples, whether BW is associated with lifespan brain changes, and to what extent lifespan effects of BW on the brain are of an environmental, rather than genetic nature. These questions, which are critical to understand how and when the human brain can be influenced through the lifespan, we address in the present study. On an overarching level, this study also addresses current debates in the field of lifespan cognitive neuroscience, namely: 1) whether consistent, reproducible relationships between phenotypes relevant for mental health and function and inter-individual differences in brain characteristics can be found ^20^, and 2) to what extent effects found in and ascribed to brain aging may actually reflect early life influences, rather than longitudinal changes in older age ^5,7,21,22^.

There are at least two different ways by which the effects of fetal growth, as indexed by birth weight, could work to produce the brain effects so far observed. 1) In line with a brain reserve model^23,24^, higher birth weight could be associated with greater brain growth before birth. This seems likely, given that the effects are seen also in young populations^6,15^. However, from the so far largely cross-sectional, or mixed models, several questions remain unanswered: Is this a fixed effect at the time of birth? Does higher birth weight also have carry-over effects to greater development in childhood and adolescence? In line with a brain maintenance model^25^, is higher birth weight associated with better maintenance of brain volumes in the face of age-related changes in older adulthood? While effects are found in young populations^5,6,15^, reduced atrophy in aging is a possible additional effect of higher BW that should be investigated, given the known relationships between birth size and brain volumes also in older age^4^. The possible effects of BW on later brain development and brain maintenance in adulthood can only be investigated by longitudinal brain imaging spanning all stages of human life. Furthermore, as BW normally reflects both genetic and prenatal environmental factors, and an environmental BW contribution to brain differences has been shown in young monozygotic (MZ) twins^15–17^, we need to study brain effects of birth weight discordance in MZ twins in this context to disentangle possible non-genetic contributions of BW through the lifespan.

We hypothesized that there are persistent effects of BW on brain characteristics through the lifespan, and hence, that these would be consistent within and across samples of varying age and origin. We test this in a Norwegian sample covering the lifespan (LCBC)^5,21^, the US developmental sample ABCD^26,27^, and the older adult UK biobank (UKB)^28,29^ sample. The associations of BW and cortical surface, thickness, volume and their change were investigated vertex-wise in a total of 5794 persons (of whom 5718 with repeated scans, and 386 monozygotic twins) with 12088 longitudinal observations, 4-82 years of age at baseline, followed for up to ∼ 8.3 years. Based on previous results^5,6,15^, we hypothesized such effects to be driven primarily by positive associations between birth weight and cortical area, with lesser, if any, effects on cortical thickness. We expected positive effects on cortical volume corresponding to positive effects on cortical area. We hypothesized that effects would be stable, so that BW mainly affects the brain “intercept” and does not relate much to brain changes. That is, we hypothesize a threshold model, whereby higher BW yields greater cortical area, and hence cortical volume, to begin with, rather than a maintenance model, whereby higher BW serves to protect against atrophy in aging. Moreover, we hypothesized that effects could not be explained solely by genetics, so that birth weight discordance in a subsample of MZ twins would also result in differences in brain characteristics through the lifespan.

## Results

Cortical surfaces were reconstructed from T1-weighted anatomical MRIs by use of FreeSurfer v6.0 (LCBC and UKB), and 7.1. (ABCD) (https://surfer.nmr.mgh.harvard.edu/)^30–33^, yielding maps of cortical area, thickness and volume. Vertex-wise analyses were run with spatiotemporal linear mixed effects modeling (FreeSurfer v6.0.0 ST-MLE package), to assess regional variation in the relationships between birth weight and cortical structure and its change. All analyses were run with baseline age, sex, scanner site, and time (scan interval) as covariates. For ABCD specifically, ethnicity was also included as a covariate. For consistency of multiple comparison corrections across analyses, the results were thresholded at a cluster-forming threshold of 2.0, p < .01, with a cluster-wise probability of p < .0.25 (p <.05/2 hemispheres).

### The lifespan relationship of BW and cortical volume, surface area and thickness

Associations of birth weight and cortical characteristics are shown in Figure 1 (for the right hemisphere, and in Supplementary Figure 1 for both hemispheres). Across all cohorts, widespread positive associations were observed between BW and cortical area. These were highly consistent across lifespan (LCBC), developmental (ABCD) and aging (UKB) cohorts, and there were bilateral overlapping effects across most of the cortical mantle. As expected, BW had in general lesser effects on cortical thickness, and no significant effects on thickness were observed in the UKB. There were however some lateral positive and medial negative effects in the LCBC and ABCD cohorts. We note that corresponding effects with increased medial frontal and occipital cortical thickness have been found associated with white matter alterations (reduced FA) in young adults born preterm with very low BW compared to term-born controls ^34^. BW was significantly positively associated with cortical volume across much of the cortical mantle. In sum, broad, bilateral, positive associations were observed across cohorts for cortical area and volume.

**Figure 1.**
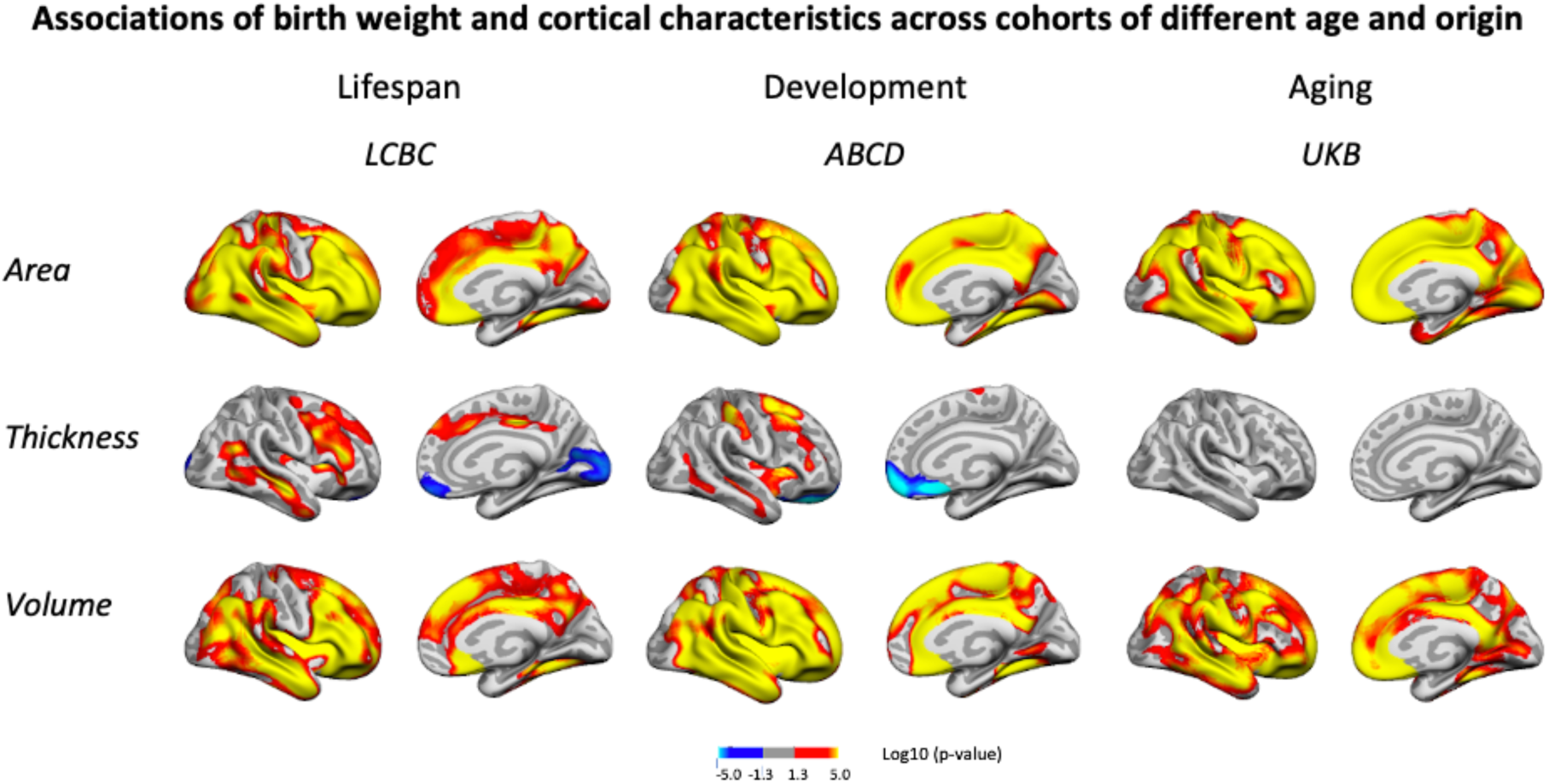
Relationships of birthweight and cortical characteristics across LCBC, ABCD, and UKB samples. Age, sex, time (interval since baseline) and scanner site (as well as ethnicity in the ABCD) were controlled for. Significant relationships are shown for area, thickness, and volume for each sample, from left to right: lateral view and medial view, right hemisphere.

Additionally controlling for education level had little effect on results (see Supplementary Figure 2). Information on gestational length (i.e. whether there was premature birth) was not available for all participants. Importantly, this information was lacking for the older participants, i.e. this information is not available for UKB, and since this information for the LCBC was drawn from the Medical Birth Registry of Norway (MBRN), only established in 1967, this was not available for the older part of the LCBC sample either. The majority of the LCBC sample and the ABCD sample, had information on gestational length, however (LCBC: n = 514; gestational length in weeks: M = 40.0 weeks, SD = 1.9, range = 25-44; ABCD: n = 3306; weeks premature: M = 1.0 weeks premature, SD = 2.1, range = 0.0-13.0). Controlling for gestational age in these subsamples had relatively little effect on results (see Supplementary Figure 3). However, as expected with reduced power in the LCBC sample, the effects in this analysis were somewhat narrower. Effects in ABCD, where almost all participants were retained for analysis, showed no sign of decrease with control for gestational length. When restricting all samples to participants with BW between 2.5 and 5.0 kg, results were also very similar (see Supplementary Figure 4). As expected from the widespread effects on cortical area and volume, effects were partly generic, with analyses controlling for ICV showing more restricted effects (see Supplementary Figure 5). However, consistent significant positive effects of BW on cortical area also when controlling for ICV were observed across all three cohorts in lateral temporal and frontal areas (see Supplementary Figure 6).

### Birth weight effects on cortical change

To test the effect of birth weight on cortical change we reran the analyses with BW x time and age x time interactions. Note BW x time (i.e., within-subject follow-up time) represents the contrasts of interest while age – and age interactions – are used to account for differences in age across individuals. Significant BW x time interactions on cortical characteristics were observed in restricted and non-overlapping regions across samples, see Figure 2 (depicting right hemisphere results, for visualization of effects in both hemispheres, see Supplementary Figure 6). Per direction of effect, the effect of BW differences was apparently reduced over time for area in LCBC and ABCD, whereas no interaction effects on area were significant in UKB. A mixture of positive (ABCD) and negative (LCBC, UKB) interaction effects were significant for thickness and volume.

**Figure 2.**
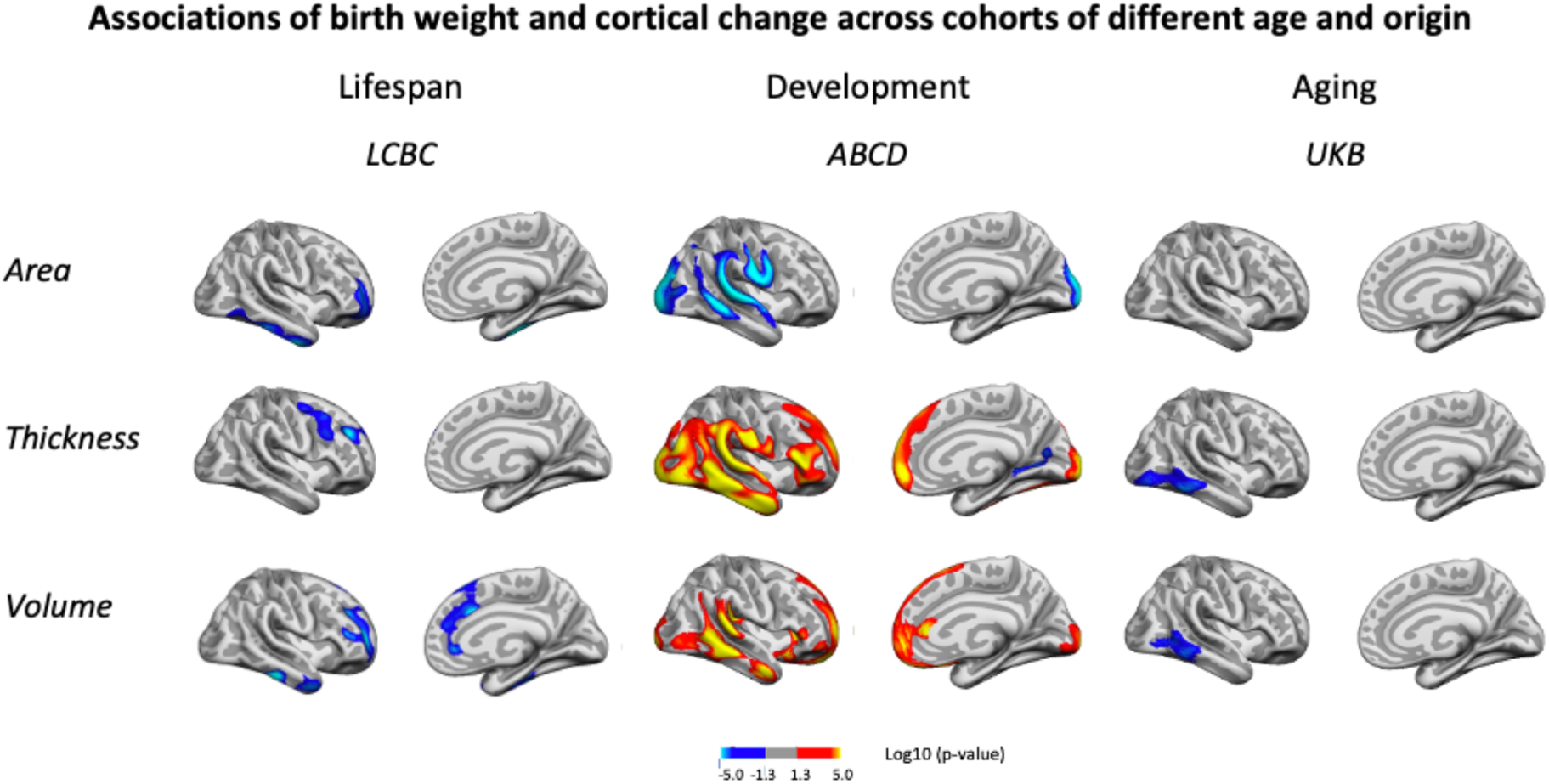
Interactions of BW and time on cortical characteristics across LCBC, ABCD, and UKB samples. Age, sex, scanner site, time, and birth weight (as well as ethnicity in the ABCD) were controlled for. Significant relationships are shown, from left to right: lateral view, right and left hemisphere, and medial view, right and left hemisphere.

Visualization of the interaction effects as seen in Figure 2 and Supplementary Figure 6, by splitting the sample in two based on BW, did not yield convincing evidence for these interactions, as shown in Supplementary Figures 7-9. In plots of LCBC data, where number of follow-ups varied, and a select portion had longer follow-up, it appeared that the effect of BW was reduced over time in these restricted regions. However, virtually parallel trajectories for the ABCD and UKB subsamples with lower and higher BW, suggested the effect size even within the areas of significant interactions of BW and time was negligible. Since UKB and ABCD samples here consisted of samples having two time points only, whereas LCBC consisted of a mix of number of follow-ups over a longer time period, there might be sample-specific selection effects also regarding other characteristics than BW that can influence these effects in LCBC. For instance, participants who do not drop out tend to have better health, cognitive ability and education, which again may relate positively to the brain measures studied here ^35,36^. Thus, caution is advised in interpreting effects seen only with longer follow-up in the LCBC sample.

Both birth weight effects on cortical characteristics and cortical change were rerun (ROI-wise) using spline models that accounted for possible non-linear effects of age on cortical structure. The results were comparable to those reported above in Figures 1 and 2. See Supplementary Figures 13 and 14 for birth weight effects on cortical characteristics and cortical change, respectively.

### Consistency of spatial relationships across and within samples

Next, we assessed whether the cortical correlates of BW (βeta-maps) showed a similar topographic pattern across the three independent datasets (UKB, ABCD and LCBC). The results showed that all the spatial comparisons were statistically significant (p < 0.05, FDR-corrected). That is, the topography of the effects of BW on cortical structure was comparable across datasets – the pairwise spatial correlation of a given cortical correlate of BW (e.g. BW effects on cortical area) was similar when estimated from two different datasets. The spatial correlations were highest for the volume measures (r = .64 -.79), and overall also high (r = .51-.71) for area measures, whereas for cortical thickness, they were more moderate (r = .24 -.45).

See spatial correlations for the right hemisphere cortical volume in Figure 3 and the full model summary in Supplementary Table 2. The results are qualitatively comparable when using -log_10_ (p) significance values instead of βeta estimates, as shown in Supplementary Table 2. The same pattern of results was largely seen also for spatial correlation of the maps capturing BW-associated cortical characteristics when controlling for ICV. The correlations were then on average somewhat lower, but there were still only significant positive correlations across LCBC, ABCD, and UKB (see Supplementary Table 2).

**Figure 3.**
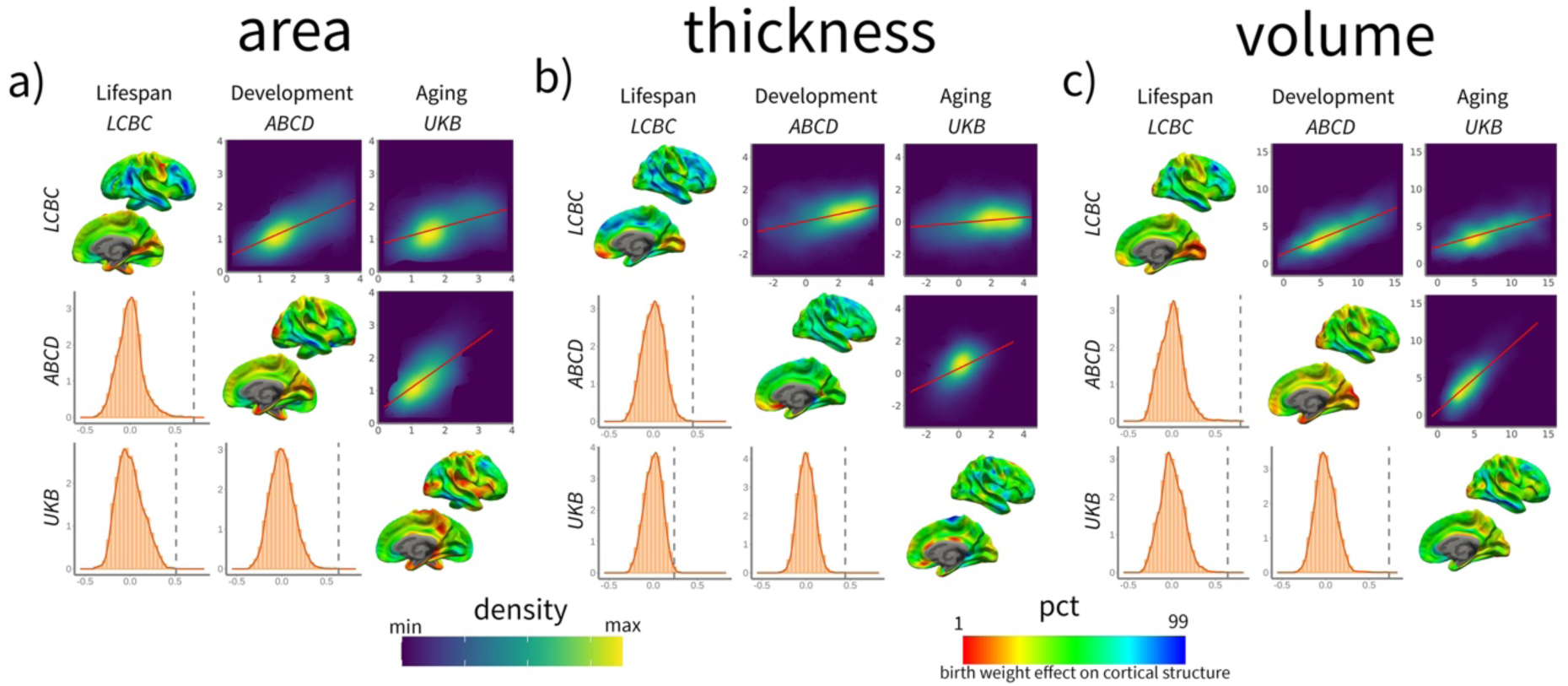
Spatial correlation of birth weight effects on brain structure across datasets for cortical a) area, b) thickness, and c) volume. Spatial correlation of birth weight effects on brain structure across datasets. For each panel, the upper triangular matrix shows Pearson’s (r) pairwise spatial correlation between the different cohorts’ cortical maps. Data is shown as a color-density plot. The red line represents the fitting between the two maps. The lower triangular matrix shows the significance testing. The dashed-grey line shows the empirical correlation, while the orange histogram represents the null distribution based on the spin test. The diagonal shows the effect of birth weight on cortical structure (right hemisphere shown only). Note that the βeta-maps are shown as a percentile red-green-blue scale, where red represents a lower (or more negative) effect of birth weight on cortical structure and vice versa. See Supplementary Table 2 for stats. Units in the density maps represent birth weight effects as mm/g, mm^2^/g, and mm^3^/g (10e^-5^) for cortical thickness, area, and volume, respectively.

In contrast, the spatial correlation of the maps capturing BW-associated cortical *change* (i.e., BW x time contrast) were either unrelated (n = 7) or showed negative associations between cohorts (n = 2). The spatial correlations of birth weight on cortical change were r = -35- xs-.05 for area, r = -.35 -.08 for volume, and r = -.20, - - .04 for thickness. See a visual representation in Supplementary Figure 10 and full stats in Supplementary Table 2.

In sum, the spatial correlation analyses imply that the different datasets show a comparable topography of BW effects across the cortical mantle – i.e. the areas more and less affected by BW were common across datasets. Thus, the BW effects on cortical structure are robust and replicable across very different datasets. In contrast, the effects of BW on cortical change are not robust across datasets, showing dissimilar topographies.

Additionally, we performed replicability analyses both across and within samples to further investigate the robustness of the effects of birth weight on cortical characteristics and cortical change. Split-half analyses within datasets were performed, to investigate the replicability of significant effects ^37,38^ of BW on cortical characteristics within samples (refer to Figure 1). These analyses further confirmed that the significant effects were largely replicable for volume and area, but not for thickness (see Supplementary Figure 11). Split-half analyses of BW on cortical change (refer to Figure 2) showed, in general, a very low degree of replicability on the three different cortical measures. See Supplementary Table 3. Replicability across datasets showed a similar pattern, that is, replicability was high for the effect of brain weight on cortical characteristics but very low for the effects of cortical change. See Supplementary Table 4 for stats. See Supplementary statistical methods for a full description of the analyses. These analyses provide complementary evidence of robust associations of BW with cortical area and volume – but not cortical change - across and within samples.

### Effects of BW discordance on brain characteristics and changes in monozygotic twins

BW discordance analyses on twins specifically were run as described for the main analyses above, with the exception that twin scans were reconstructed using FS v6.0.1. for ABCD and the addition of the twin’s mean birth weight as a covariate. BW discordance was associated with cortical area, where the heavier twins had greater area in some frontal, temporal and occipitotemporal regions, with effects in the right hemisphere only surviving corrections for multiple comparisons. We note that these regions mostly overlap with regions where positive effects of BW were also seen in the bigger sample. Strikingly, the effect of BW discordance, as shown in Figure 4, appeared similar in size to the effect of BW itself in the MZ twin sample. However, note that this plot is merely for illustrating effects, the effect size is inflated for the BW discordance plot, since the values are derived from areas already identified as significantly related to BW. There was no association of BW discordance and cortical area changes over time.

**Figure 4.**
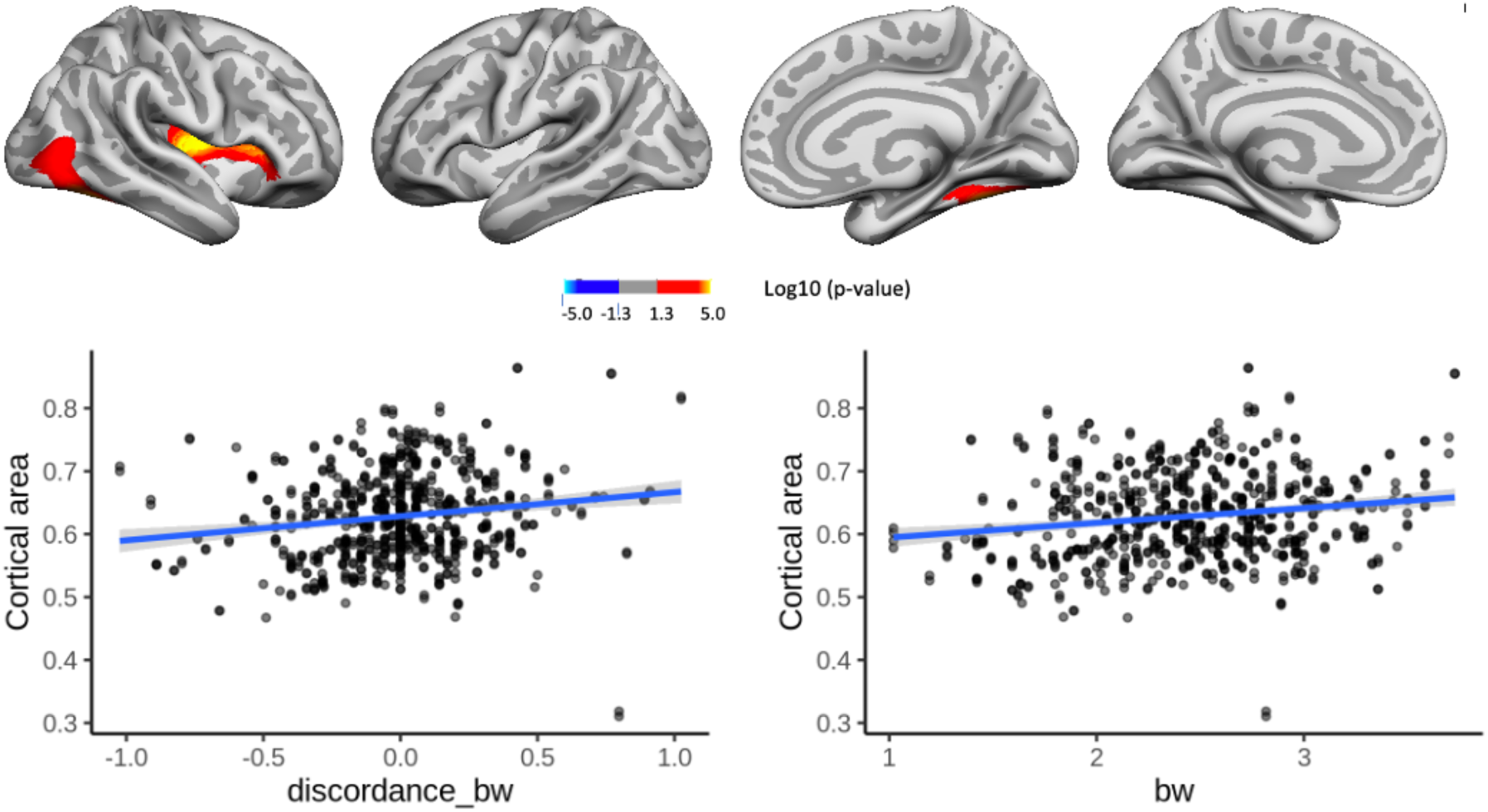
Effects of birthweight discordance on cortical area in the sample of monozygotic (MZ) twins. Significant relationships are shown from left to right: lateral view, right and left hemisphere, and medial view, right and left hemisphere. Plots are showing -for illustrative purposes – individual data points and expected trajectories for cortical area in mm (Y-axes) within the significant regions according to birth weight (BW) discordance (left panel) and BW (right panel) in kilograms (X-axes).

BW discordance also had a significant negative effect on cortical thickness in restricted right frontotemporal regions, where being the lighter twin yielded greater thickness. These significant effects did not appear to overlap with regions where significant negative associations with BW were seen in the bigger sample. BW had little effect on cortical thickness in the significant region, and the effect of BW discordance in the identified regions, as shown in Figure 5, appeared greater than the effect of BW itself here in the MZ twin sample. However, this plot is merely for illustrating effects, it should be noted that the effect size is inflated for the BW discordance plot, since the values are derived from areas already identified as significantly related to BW.

**Figure 5.**
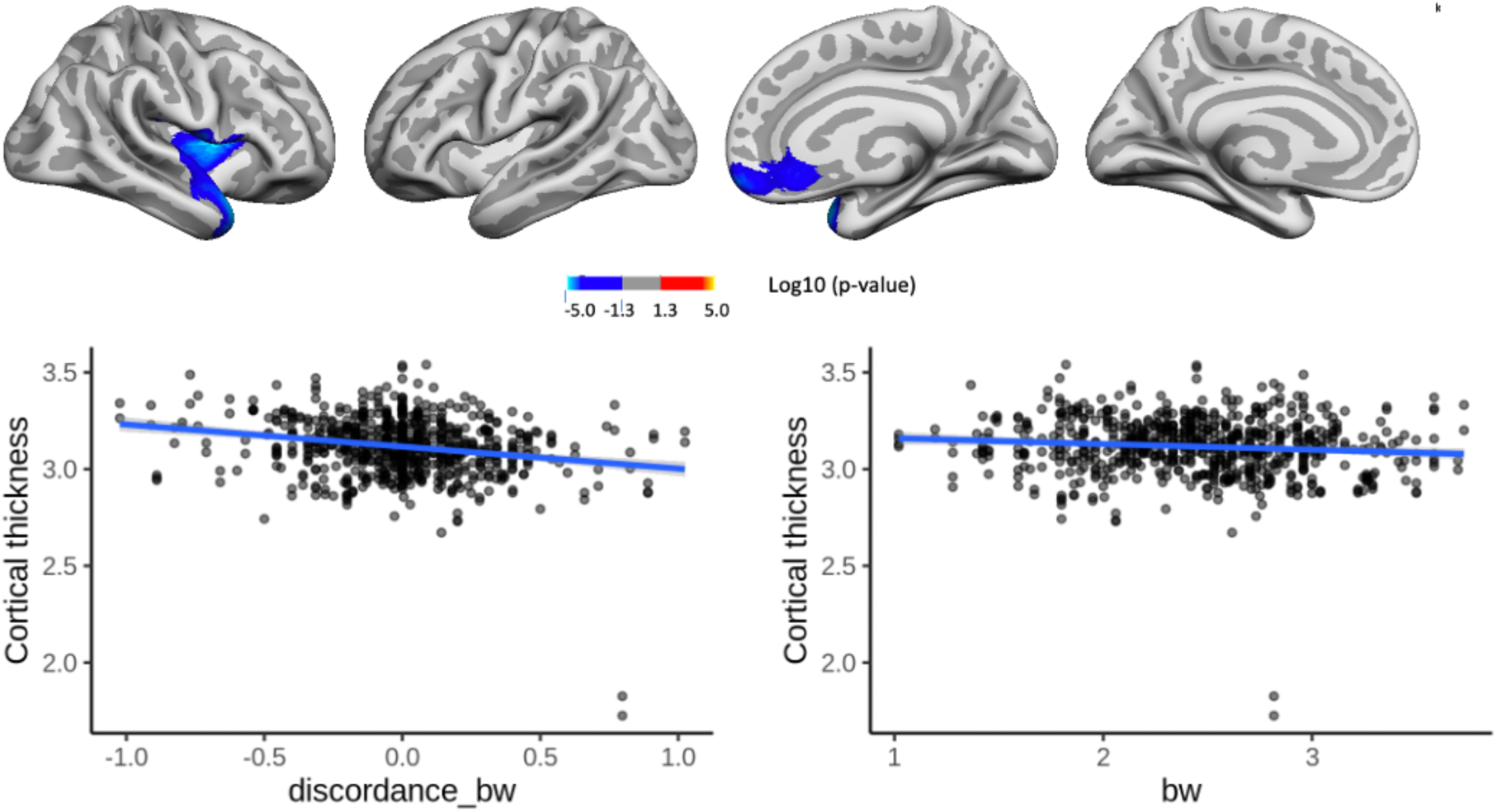
Effects of birthweight discordance on cortical thickness in the sample of monozygotic (MZ) twins. Relationships significant corrected with cluster-forming threshold of 2.0 (p< .01) are shown from left to right: lateral view, right and left hemisphere, and medial view, right and left hemisphere. Plots are showing – for illustrative purposes - individual data points and expected trajectories for cortical thickness in mm (Y-axes) within the significant regions according to birth weight (BW) discordance (left panel) and BW (right panel) in kilograms (X-axes).

In a very small area of the right hemisphere, there was a significant association of BW discordance and cortical thickness change, meaning the lighter twin had greater cortical thickness over time, but this effect was both regionally and quantitatively minor, as shown in Supplementary Figure 12. There were no significant effects of BW discordance on cortical volume or volume change over time.

Finally, to formally assess whether the cortical correlates (βeta-maps) of BW discordance in the twin subsample corresponded to cortical correlates of BW in the bigger samples, we did a meta-analysis of these estimates for area, thickness and volume in the UKB, ABCD and LCBC, and then assessed whether the cortical correlates of BW and BW discordance (βeta-maps) showed a similar topographic pattern across the datasets. The results of this meta-analysis-twin comparison showed only positive relationships, for area, r = .23, thickness r= .19, and volume r =.22. However, the respective uncorrected p-values were .08, .12, and .04, so the spatial comparisons would not be statistically significant (p < 0.05, FDR-corrected). However, the positive correlations are suggestive that the topography of the effects of BW discordance in genetically identical twins on cortical structure was to some extent comparable to effects of individual differences in BW in the bigger sample.

### Effects of BW differences relative to other estimated effects in aging

We calculated the effect of 1 SD difference in BW (on average 600 g, see Table 1) on cortical and brain volume across cohorts, to illustrate what variance is captured here by the early life effects, relative to the estimates of later aging changes. The effect of 1SD lower BW on cortical volume was 6708 mm^3^, 8466 mm^3^, and 5980 mm^3^ in LCBC, UKB and ABCD, respectively. This was equal to 1.2%, 1.6% and 1.1% lesser cortex with 600-700 grams lower BW for each sample, respectively. In the context of brain aging, this would be a substantial effect, higher than most risk/protective factors for dementia outlined by the Lancet Commission on dementia prevention ^39,40^ which include smoking, education, obesity, and alcohol amongst others (see^22^ for a summary of the different effect sizes). The estimated yearly cortical volume reduction from 50 to 60 years is 895 mm^3^ in the LCBC and 1402 mm3 in the UKB samples, respectively. Hence, the effect of 600 grams difference in BW could, if only cross-sectional data were available, be quantitatively equal to that of 7.5 and 6 years of estimated age differences in LCBC and UKB, respectively. Many factors that influence cortical volume have nothing to do with either age or differences in general brain function, including e.g. sex, so this analogy we only include to illustrate what might happen if such variance were to be ascribed to age, e.g. as in modeling “brain-age”^7^, see further discussion below.

**Table 1.** Descriptive statistics for the longitudinal samples. F = number of females in the sample, M = Mean, SD= Standard deviation. Numbers are given in years for baseline age, time since baseline and education, birthweight is given in kilograms. For LCBC, only 584 participants had information on education. Parental education was used in ABCD, and in LCBC when the participant was below 18 years of age, and also if no other education information was available for participants up to 21 years.

| Study | N | F | Scan<br>s | Baseline age |  |  | Time since<br>baseline |  |  | Birth weight |  |  | Education |  |  |
| --- | --- | --- | --- | --- | --- | --- | --- | --- | --- | --- | --- | --- | --- | --- | --- |
|  |  |  |  | M | SD | Range | M | SD | Range | M | SD | Range | M | SD | Range |
| LCBC | 635 | 350 | 1922 | 19.1 | 20.7 | 4.1-81.9 | 2.6 | 2.4 | 0.1-8.3 | 3.5 | 0.6 | 0.9-6.0 | 17.1 | 2.4 | 9-24 |
| ABCD | 3324 | 1562 | 6648 | 10.0 | 0.6 | 8.9-11.1 | 2.0 | 0.1 | 1.4-2.8 | 3.2 | 0.7 | 1.0-6.7 | 16.0 | 2.6 | 6-21 |
| UKB | 1759 | 1009 | 3518 | 62.0 | 7.1 | 47-80.3 | 2.3 | 0.1 | 2.0-2.9 | 3.4 | 0.6 | 0.9-6.4 | 14.2 | 2.4 | 7-16 |

## Discussion

The present results indicate that BW, the earliest widely and easily obtainable congenital metric, show robust, persistent, and chiefly stable associations with brain characteristics through life. Especially, BW was associated with cortical area and volume in an age and time-invariant fashion. The robustness of this effect is quite remarkable, given the wealth of different influences individuals meet after birth, which are repeatedly assumed and reported to have major impact on the brain through the protracted human lifespan ^39,40^. It is also in quantitative terms outstanding, compared to consistency of cortical topographies reported for other phenotypical factors ^20^. This is also special for a phenotype known to be environmentally influenced, unlike biologically hardcoded phenotypes such as sex or age, for which there are known brain-wide association studies (BWAS) patterns ^37,41^.

Typically, other factors relating to later socioeconomic status, lifestyle, and health, get the most attention in adult and aging brain research ^39,40^. Such factors, which are then targeted for prevention and intervention at different stages of the life-course, often do not show consistent relationships to brain characteristics ^42^, may not actually be causal ^42^, and may themselves be related to prenatal growth ^19^. Another phenotype which obviously, like BW, reflects both genetic and environmental influences ^12–14^, is body mass index (BMI) ^43^. Consistent BWAS patterns have been reported for BMI ^37^. BW stands out as the single chronological earliest phenotype, and besides BMI, BW appears to have the most replicable and consistent relations to cortical morphology, as shown here both across and within samples. It has been claimed that smaller than expected brain–phenotype associations and variability across population subsamples can explain widespread replication failures for brain-wide association studies (BWAS) ^20^. However, this is necessarily a question of which phenotypes are the most relevant to relate to brain characteristics. Also, the temporal order of factors needs to be considered if causal interpretations are to be made. Chronologically later factors necessarily do not cause earlier ones. While we cannot claim that BW itself causes the cortical characteristics observed in aging, the cortical variance explained by BW after one decade, and seven or eight decades of life alike, is unlikely to be explained by influences only present at some point in adulthood or aging. BW, as further discussed below, depends on genetic, as well as prenatal environmental influences ^12–14^, which likely have causal effects on early brain morphological features. Here we find that these effects are substantial also in the aging brain.

We calculated that a BW difference of one SD (about 600 g) equaled a difference in cortical volume on the order of 1.1-1.6 percent in these cohorts. This is a quite big effect, of a magnitude relevant for explaining a substantial portion of the differences typically seen between patients with neurodevelopmental or neurodegenerative diseases and healthy controls. As noted, BW differences have been reported for neurodevelopmental disorders such as ADHD ^44^, and also other neurodevelopmental disorders, such as schizophrenia or more general psychopathology, with the cortical effects detected not invariably being very large in absolute terms ^45^. There is a limit to the range of variation that can apply to human cortical volumes, in general, (e.g. virtually none have cortical volumes below 0.45 or above 0.65 l). In terms of sample representativity, one may assume that there can be a restriction relative to the actual range of human cortical volume variation, as the present samples specifically are largely healthy ^46^. Much of the differences within this limited range of variation are explained by factors we controlled for, such as age and sex. With the present effect, on the restricted range of variation, combined with big samples, it is obvious that BW differences of much less than our example magnitude (600 g) may be detectable in the cortical morphology of patients versus controls. In the context of aging and neurodegenerative change, the estimated cortical effect of ≈600 g difference in BW is of a magnitude many times the annual cortical reduction estimated to take place from e.g. 50 to 60 years in the adult cohorts. This is a substantial effect in brain imaging and may illuminate why metrics such as “brain age”, assumed to index aging-related processes, may rather largely capture variance already determined at birth ^7,47,48^. Neglecting this especially consistent and early factor is likely to lead to a substantial portion of human brain variance being either erroneously ascribed to factors only present at later life stages ^7,22^ or left unaccounted for.

The solidity, replicability, and universality of effects as shown here for a partly environmentally influenced metric ^12–14^, appear exceptional in human brain imaging. The within-sample replicability results are not fully comparable to other studies assessing the replicability of brain-phenotype associations due to analytical differences (e.g. sample size, multiple-comparison correction method) ^20,37^. Still, these results too clearly show that the rate of replicability of BW associations with cortical area and volume are comparable to benchmark brain-phenotype associations such as age and BMI with brain structure ^37^. The BW-cortical volume and area associations may be among the topographically broadest and most consistent effects so far seen as stable across the lifespan of the human brain. The three cohorts studies differ on a range features known to be highly and reliably related to cortical characteristics, first and foremost age ^2,5,37,49,50^, but also country of origin and representativity of the populations from which they are drawn ^27,51^. Yet there is a comparable topography of BW effects across the samples. This is so despite the samples collectively spanning the entire human age range, within which there are always substantial age-related changes in cortical structure ^2,5,52–54^. The present results thus indicate that fetal growth influences an offset of brain reserve ^23,24^ and that this brain reserve effect is persistent and stable through the lifespan.

In contrast, the cortical maps capturing *change* in cortical structure associated with BW were not robust across datasets; i.e. the most positive and negative association with BW on cortical change did not overlap at all between the different cohorts. While there was evidence from ABCD that BW affected regional cortical development in the narrow age range covered, there were limited and no consistent effects of BW on cortical change across cohorts. Importantly, there was no indication whatsoever that BW could be associated with better brain maintenance ^25^ in the face of age-related changes in older adulthood. Thus, the data seem to indicate that any effect of BW on cortical *change* may be of relatively more temporary nature. The “offset effect” of BW, on the other hand, appears persistent and consistent, especially in terms of stable and widespread effects on cortical area and volume across the lifespan.

The sensitivity analyses indicate that the associations between BW and cortical characteristics are seen irrespective of not only sex and age, but also education, head size (ICV), and cases of abnormal BW. Such patterns could point to an underlying genetic pleiotropy of BW and brain characteristics. Interestingly, however, recent findings indicate that effects of exposure to environmental adversity on epigenetic programing in aging may be localized to the in-utero period^55^. The effects of BW discordance in MZ twins in this context, align with other studies^16,17^ pointing to also non-genetic, that is environmental, influences in the womb, associated with the pattern observed for cortical area effects. These analyses also account for multiple possibly confounding variables that could represent a mix of genetic-environmental effects, such as parental socioeconomic status, parity, or prenatal exposures shared between twins in the same womb such as maternal smoking or use of alcohol.

The neural basis for the observed association cannot readily be ascertained from human imaging studies tracking change^5,6,15^. While the “fetal origins hypothesis”, proposing that cardiovascular disease in adulthood is related to undernourishment in utero^56,57^ is well-known, there has been focus on “brain-sparing” adaptations under such conditions ^19^. However, our finding that early human development in utero appears to be associated with a persistent and stable brain reserve effect, is largely in correspondence with what is known of human nervous system development and change through the lifespan: While synaptogenesis, synaptic remodeling and myelination are known to be protracted processes long after infancy^58–61^, numerous processes in brain development appear to be exclusively, or almost exclusively happening before birth. For instance, neurogenesis takes place almost only in fetal development ^62^. Even if controversies remain, evidence suggests that any adult human neurogenesis must be severely restricted in location and amount^63,64^. Thus, human beings appear to be born with almost all cortical neurons they will have through life, and neuronal migration and differentiation are also defined early, by the place and time the neuron is born during fetal life^62^. Factors that affect placental function and uterine and/or umbilical blood flow on a chronic basis may lead to restricted fetal growth, including brain growth, and given the timing of brain development, it may not be surprising that effects would be stable across years. Animal studies of chronic placental insufficiency have shown effects on brain development which persist with age^65^. Hence, the relationship between BW and cortical characteristics in the normal population could likely have a twofold etiology: it is likely to in part be based on normal variation in genetically determined body and brain size, but it also may be based on variations in environmental prenatal conditions, yielding differences in optimality of early brain development persisting through the lifespan.

These results indicate that there is potential to increase brain reserve throughout the lifespan, also in aging, by combating factors affecting fetal growth negatively. While it is unknown to what extent results from these specific US and European samples can be generalized to other populations, the current potential to improve prenatal factors may be especially high in low and middle-income countries, where the demographic changes will also be more marked in terms of the aging population^66^. About 200 million children in developing countries are not meeting their growth potential, and improving the prenatal environment is likely important to help children reach their full potential^67^ – and ultimately also to help them stay above a functional threshold into older age. Also in industrialized countries, including the US, environmental factors are associated with birth weight. The opioid epidemic is increasingly affecting pregnant women, and in-utero opioid exposure is associated with higher risk of fetal growth restriction^68^. Among highly common exposures, air pollution may for instance be of relevance. Recently, local traffic congestion-pollution exposure during pregnancy, independently of a series of maternal sociodemographic characteristics, was associated with reductions in term BW in a large US sample^14^. Programs and policies to limit such environmental factors reducing fetal growth may thus enhance brain reserve and ultimately prevent more people from falling below a functional threshold even in advanced age.

### Limitations

Some limitations should be noted. First, for most of the participants, only self-reported or parent-reported BW was available. While there was a very high correlation between registry and self-reported BW in LCBC, this is a possible source of noise. Second, pregnancy-related information of possible relevance, such as gestational age at birth, complications, method of delivery, maternal disorders, smoking, alcohol and drug intake, was not available across all participants of the different samples, and was thus not analyzed here, or, as for gestational age at birth, could only in part be controlled for. Some of these factors may be systematically related to BW, and may thus represent confounds^69^. There were some premature, and very low BW participants in the samples, and these conditions are associated with known reductions in cortical volume^70,71^. However, the analyses controlling for gestational age, as well as on the restricted range of BW – excluding very preterm and very/extremely low BW children – and the analysis controlling for education, which may again relate to some of these factors, showed very similar results. It is unknown to what extent the BW of participants reflect their individual fetal growth potential, as a fetus with normal BW can be growth restricted and a fetus with low BW can have appropriate growth^72^. We believe, however, that possible differences in such factors would likely serve to decrease consistency of results, and not lead to inflated estimates of consistency.

Finally, it is beyond the scope of the present study to relate BW and cortical characteristics through the lifespan with cognitive functional differences. While all the cohorts included here also have some measures of cognitive function, they vary across samples. Furthermore, tests of cognitive function are, relative to brain imaging metrics, much more prone to test-specific test-retest effects. Thus, assessing the stability of effects across cohorts would be challenging. We note that, e.g. in twins, even though there are data to suggest a relationship between BW differences and neuroanatomical features^15–17^, and BW differences and differences in cognitive function^73^, twins discordant for BW and neuroanatomical features may not show significant differences in neurodevelopmental outcomes^17^. There are many factors that influence cortical volume that have nothing to do with either age or differences in general brain function, including e.g. sex or overall differences in height. We cannot from the present data draw conclusions about effects on individual differences in cognitive function or its change across the lifespan. Indeed, part of birth weight effects on brain structure can be explained by overall somatic growth. This study does not provide one specific mechanism by which BW is associated with brain structure. Indeed variance explained by different mechanisms may vary across samples. Further studies are needed to illuminate such questions, the present study is primarily designed to answer the question of whether associations of BW and cortical structure are consistently found throughout the lifespan.

## Conclusion

The current results show that a simple congenital marker of early developmental growth, BW, is consistently associated with lifespan brain characteristics. While some significant effects of BW on cortical change patterns were also observed, these were regionally smaller and showed no consistency across cohorts. In conclusion, while greater early human developmental growth does not appear to promote brain maintenance in aging, it does, in terms of greater cortical volume and area, relate positively to brain reserve through the lifespan. Thus, there appears to be an omnipresence of fetal factors in the spacetime of the human brain through the lifespan. This indicates a potential to increase brain reserve at all ages, including in aging, by combating factors affecting fetal growth negatively. Given the exceptional consistency and broadness of this cortical topographical effect, it should be taken into account in studies of brain research on individual differences, whether the brains studied are those of eight- or eighty-year-olds.

## Materials and methods

### Samples

In total, longitudinal data for 5718 persons with 12088 MRI scans from the LCBC, ABCD and UKB studies were included in the analyses. For UKB the dataset released February 2020 was used. For ABCD, the Data Release 3.0 was used (s<u>ee</u> http://dx.doi.org/10.15154/1528313 for this NDA study). Only persons with longitudinal MRI scans were included in the main analyses, to limit the possibility that estimates of change were biased by immediate sample selection effects (i.e. those that remain for follow-up are known to have other characteristics than those who have only one time-point assessment in longitudinal studies, and this can bias effects). However, for the separate MZ twin-analyses, we also included participants with only one time-point MRI, to obtain an age-varying sample for assessment of whether non-genetic effects were found throughout the lifespan, including in adulthood and older age. Of the 386 MZ twins included, 310 had longitudinal imaging data. The twins were mostly (n = 310) from the developmental ABCD sample (age 10-11), whereas 64 adults were from LCBC (age 18-79 years), and 12 were from UKB (age 50-80 years). All samples consisted of community-dwelling participants. For the most part, these were recruited by means of some type of population registry information (see Supplementary Materials, SM), but part of the LCBC cohort consisted of convenience samples, i.e. the studies were advertised broadly. Part of the LCBC sample was recruited through the Norwegian Mother, Father and Child Cohort Study (MoBa)^74^, and the Norwegian Twin Registry (NTR)^75^. Thus, this study includes data from MoBa and NTR, and both studies are conducted by the Norwegian Institute of Public Health. LCBC participants were part of observational studies, but subsamples were part of studies including cognitive training (n = 168). As BW was not a criterion for assigning participants to cognitive training, these were included here. For the majority of participants, and all in UKB and ABCD, BW was collected as self-report or, for children, parent report, at the time of scan. For LCBC, BW was for the majority (n = 526 collected from the Medical Birth Registry (MBRN), available for those recruited through MoBa, or NTR, or if collected by consent for participants born 1967 and later, and in part by self-report in connection with scanning, or earlier self-report to the Norwegian Twin Registry (for twins recruited through this registry). MBRN is a national health registry containing information about all births in Norway. Comparative analyses in the LCBC sample for 354 persons who had available both MBRN records and self-report/parent-report of BW, showed a very high correlation of BW as obtained from the different sources (r =.99). A high reliability of self-reported BW over time has also been found in broader NTR samples ^76^ Demographics of the samples in the main analyses are given in Table 1, see SM for details.

### Statistical Analyses

#### Cortical Vertex-wise Analyses

Reconstructed cortical surfaces were smoothed with a Gaussian kernel of 15 mm full-width at half-maximum. We ran vertex-wise analyses to assess regional variation in the relationships between birth weight and cortical structure; area, thickness and volume at baseline and longitudinally. In all models we included 1) baseline age, sex and scanner site, as well as time (scan interval) as covariates. For ABCD specifically, ethnicity was also included as a covariate, as this sample is recruited to have and has, ethnic variation (see Supplemental Methods for details), whereas the other samples entered in the present analyses had little ethnic variation (i.e. in UKB, >98% of participants included in the present sample defined themselves as British/Irish/Any other white background. In LCBC, this information was unfortunately not encoded for all, but the sample was mainly of white background). In further models, we additionally included education as a covariate. General linear models were run in turn using as predictors: birth weight, the interaction term birthweight x scan interval, and the interaction term of baseline age x time (scan interval) x birth weight. When analyses were run with baseline age x scan interval x birth weight as predictor, the interaction terms of baseline age x scan interval, scan interval x birth weight, and birth weight x baseline age were included as additional covariates. Standardized values were used in analyses for age, scan interval, BW, BW discordance, and education. For consistency of multiple comparison corrections across analyses, the results were thresholded at a cluster-forming threshold of 2.0, p < .01, with a cluster-wise probability of p < .0.25 (p <.05/2 hemispheres). Finally, models were rerun only including participants with birth weights between 2.5 and 5.0 kg, to assess whether relationships were upheld also when excluding low and high birthweights. Given previous findings of broad effects of BW on cortical area and volume ^5,6,18^, we did not expect effects to be localized. Rather, we expected BW to affect gross head and brain size irrespective of sex, but we also performed supplementary analyses controlling for intracranial volume (ICV) in order to check for possible specificity of effects. Spatial correlation analyses ^77–79^ were run on the cortical maps (for more information see SI) for analyses results using BW as predictor, from LCBC, ABCD, and UKB, to assess the overlap of BW-cortical characteristics associations in terms of topography and effect sizes. In a separate set of analyses, we restricted the sample to only monozygotic twins, and studied effects of BW discordance (number of grams BW above or below MZ twin). In these models, we included time, baseline age, sex and site as covariates.

## Abbreviations

BW: birth weight;
MRI: Magnetic Resonance Imaging

## Acknowledgements

Data used for the LCBC sample were in part obtained through the Medical Birth Registry (MBRN) of Norway, the Norwegian Twin Registry (NTR) and the Norwegian Mother, Father and Child Cohort Study (MoBa). MoBa is supported by the Norwegian Ministry of Health and Care Services and the Ministry of Education and Research. We are grateful to all the participating families in Norway who take part in this on-going cohort study. Data used in the preparation of this article were obtained from the Adolescent Brain Cognitive Development (ABCD) Stud<u>y</u> (https://abcdstudy.org), held in the NIMH Data Archive (NDA). This is a multisite, longitudinal study designed to recruit more than 10,000 children age 9-10 and follow them over 10 years into early adulthood. A listing of participating sites and a complete listing of the study investigators can be found at https://abcdstudy.org/consortium_members/. ABCD consortium investigators designed and implemented the study and/or provided data but did not necessarily participate in the analysis or writing of this report. This manuscript reflects the views of the authors and may not reflect the opinions or views of the National Institutes of Health (NIH) or ABCD consortium investigators. The ABCD data used in this report came from DOI 10.15154/1504041 and 10.15154/1520591. DOIs can be found at https://dx.doi.org/10.15154/1528313. Part of the research was conducted using the UK Biobank resource under application number 32048. The ERC, RCN, EC, NIH and UK Biobank had no role in the design and conduct of this specific study. We thank all participants for contributing to all the respective data sources.

This study was funded by ERC grants (771375 and 313440 to KBW; 283634 and 725025 to AMF), Research Council of Norway (RCN) Grants (to KBW, AMF and DVP), and European Commission (EC) EU Horizon 2020 Grant agreement number 732592 (Lifebrain). The ABCD Study® is supported by the National Institutes of Health and additional federal partners under award numbers U01DA041048, U01DA050989, U01DA051016, U01DA041022, U01DA051018, U01DA051037, U01DA050987, U01DA041174, U01DA041106, U01DA041117, U01DA041028, U01DA041134, U01DA050988, U01DA051039, U01DA041156, U01DA041025, U01DA041120, U01DA051038, U01DA041148, U01DA041093, U01DA041089, U24DA041123, U24DA041147. A full list of supporters is available at https://abcdstudy.org/federal-partners.html.

## Data Availability

The code used for vertex-wise ST-LME analyses and spatial correlations analyses can be accessed here https://github.com/LCBC-UiO/paper-birthweight-brainchange-2022. Two of the datasets used, the ABCD https://abcdstudy.org and UKB https://www.ukbiobank.ac.uk databases are open to all researchers given appropriate application, please see instructions for how to gain access here: https://nda.nih.gov/abcd/ and https://www.ukbiobank.ac.uk/enable-your-research. The LCBC dataset has restricted access, requests can be made to the corresponding author, and some of the data can be made available given appropriate ethical and data protection approvals. However, the registry data on birth weight connected to this sample are not shareable by the authors, as these data are owned by the Medical Birth Registry of Norway, https://www.fhi.no/en/hn/health-registries/medical-birth-registry-of-norway/medical-birth-registry-of-norway/, the Norwegian Mother, Father, and Child Cohort Study https://www.fhi.no/en/studies/moba/, and the Norwegian Twin Registry https://www.fhi.no/en/more/health-studies/norwegian-twin-registry/ so that any access to data must be approved by them. Group-level unthresholded p-maps, F-maps, Beta-maps, and degrees of freedom for the univariate analyses accompany this manuscript as additional material.

## Supplementary Material

## Supplementary text

## Supplementary Methods

### Samples

#### LCBC

##### Population, Recruitment and General Description of Study/ Procedures

Cognitively healthy, community dwelling participants across the lifespan were drawn from studies coordinated by the Research Group for Lifespan Changes in Brain and Cognition (LCBC www.oslobrains.no), approved by a Norwegian Regional Committee for Medical and Health Research Ethics (REK South-East). Written informed consent was obtained from all adult participants and from parents or other legal guardians for participants below age of majority. The samples were recruited in part by newspaper and web page adds, and in part by population registry-based research studies. Part of the developmental sample was recruited through the population registry-based study MoBa, the Norwegian Mother, Father and Child Cohort Study at The Norwegian Institute of Public Health https://www.fhi.no/en/studies/moba/ ^80^. The Norwegian Mother, Father and Child Cohort Study (MoBa) is a population-based pregnancy cohort study conducted by the Norwegian Institute of Public Health. Participants were recruited from all over Norway from 1999-2008. The establishment of MoBa and initial data collection was based on a license from the Norwegian Data Protection Agency and approval from The Regional Committees for Medical and Health Research Ethics. The MoBa cohort is currently regulated by the Norwegian Health Registry Act. The current study was approved by The Regional Committees for Medical and Health Research Ethics (REK sør-øst C 2010/2359). Part of the adult sample was recruited through the Norwegian Twin Registry https://www.fhi.no/en/more/health-studies/norwegian-twin-registry/ ^75^. The current NTR sub-study was approved by The Regional Committees for Medical and Health Research Ethics (REK South-East C 2018/94). Both MoBa and NTR are linked to The Medical Birth Registry (MBRN), which is a national health registry containing information about all births in Norway. By individual consent, information on BW was also obtained from the MBRN also for some LCBC participants not being part of MoBa or NTR. Approval for these studies was given by The Regional Committees for Medical and Health Research Ethics (REK Sør-Øst B 2017/653). Most participants, including all children, were recruited for observational studies, while some adults were recruited to enter into cognitive training studies after baseline assessment.

##### Inclusion/Exclusion Criteria/ Screening

Adult participants were screened using a standardized health interview prior to inclusion in the study. Participants with a history of self- or parent-reported neurological or psychiatric conditions, including clinically significant stroke, serious head injury, untreated hypertension, diabetes, and use of psychoactive drugs within the last two years, were excluded. Further, participants reporting worries concerning their cognitive status, including memory function, were excluded. Participants above 40 years scored ≥26 on the Mini Mental State Examination ^81^. Additionally, a generalized additive mixed model regressing area/thickness/volume nonlinear on age with random intercept per participant, was fitted independently at each of 163842 vertices. 60 observations were defined as outliers and excluded, as the absolute value of their residual was large than four times the residual standard error at more than 6000 vertices.

##### Variables Used for Birth weight and Education

When possible, by participant consent, birth weight was obtained from the records of the Medical Birth Registry of Norway (MBRN, for participants born 1967 when the Registry was started, or after). MBRN records of birth weight was obtained for at total 526 of the participants. For 15 participants, birth weight records based on historical self-report to the Norwegian Twin Registry were obtained. Otherwise, self-report, or for children, parental report, of birth weight in connection with participation in the MRI scanning studies were used. For analyses controlling for gestational length, only information from the MBRN was used. For education, if multiple values were reported across timepoints, the highest was chosen. Education was recorded as total years of education to the highest obtained degree, for adults, and for participants < 20 years of age, average of parental education was used. However, for some participants in the age range up to 21.3 years at baseline scan, parental education was used, if no report of own education existed. Age was recorded in years and months at the time of the baseline MRI scan.

##### MRI Scanning and Processing

Brain imaging data were collected across 2 sites on 1.5 T and 3T scanners (Siemens Avanto, Skyra and Prisma (Siemens Corp., Erlanger, Germany, se Supplementary Table 1). At baseline 345 participants were scanned at Avanto 1 at Oslo University Hospital, Oslo, and 105 were scanned at Avanto 2 at St. Olav’s Hospital, Trondheim, 74 were scanned at Prisma and 111 at Skyra, Oslo University hospital. All were followed up longitudinally at least once at the same scanner. To the extent that scanners were switched for longer follow-up, double scanning (at old and new scanner) was implemented to the extent possible, and both scans were from that time point was entered into analysis. 369 participants did not switch scanner during the study, 266 switched scanner at least once, of which 106 participants were scanned with both scanners at the same time point. MRI data were processed using FreeSurfer 6.0.

#### ABCD

##### Population, Recruitment and General Description of Study/ Procedures

The primary aim of the Adolescent Brain Cognitive Development (ABCD) study https://abcdstudy.org is to track human brain development from childhood through adolescence to determine biological and environmental factors that impact or alter developmental trajectories ^26^. ABCD has recruited > 10 000 9-10 years olds across 21 US sites with harmonized measures and procedures, including imaging acquisition https://abcdstudy.org/scientists-workgroups.html. A goal of the ABCD study is that its sample should reflect, as best as possible, the sociodemographic variation of the US population ^27^. Of relevance to the present analyses, children were ineligible to participate if they had any MRI contraindications, or prematurity at birth <28 weeks ^8,82^. For ABCD, the birth weight values were extracted from the dataset release 2.0.1 at baseline (consisting of a total of 11875 participants), and the remainder of data were extracted from release 3.0.

##### Inclusion/Exclusion Criteria for present analyses

All participants who had birth weight reported and had underwent MRI scanning at more than one timepoint with FreeSurfer Quality Control OK were included in the main analyses. Additionally, a generalized additive mixed model regressing area/thickness/volume nonlinear on age with random intercept per participant, was fitted independently at each of 163842 vertices. 223 observations were defined as outliers and excluded, as the absolute value of their residual was large than four times the residual standard error at more than 6000 vertices.

##### Variables used for Birth Weight and Education

We used the fields devhx_2_birth_wt_lbs_p and devhx_2b_birth_wt_oz_p. These fields were converted to grams, and added together. If devhx_2b_birth_wt_oz_p was missing, it is set to 0, so that the only contribution came from devhx_2_birth_wt_lbs_p . If devhx_2_birth_wt_lbs_p was missing, the final birth weight value was set to NA, and hence the participant was excluded from analysis. In cases where birth weight was reported multiple times for one participant, and values deviated, participants were excluded from analysis if the discrepancy was greater than 10%, else the mean reported number was entered. For the analyses controlling for gestational length, we used the variable weeks_premature. Education was entered as the maximum education of the parents in years.

##### MRI Scanning and Processing

Brain imaging data were collected across 21 sites on 3T scanners (Siemens Prisma (Siemens Corp., Erlanger, Germany), GE Discovery MR750 (GE Healthcare, Chicago, IL), and Philips Achieva (Philips, Amsterdam, the Netherlandsfewion parameters are listed in ^26^ and at https://abcdstudy.org/images/Protocol_Imaging_Sequences.pdf. Images were processed using the FreeSurfer 7.1.0 software package for the analysis of the man longitudinal sample, whereas for the MZ tiwn analysis (done collectively withUKB and LCBC MZ samples), the FreeSurfer 6.0 software package was used.

#### UKB

##### Population, Recruitment and General Description of Study/Procedures

UK Biobank (UKB) (https://www.ukbiobank.ac.uk/about-biobank-uk/) is a major national and international health resource with the aim of improving the prevention, diagnosis and treatment of a wide range of illnesses. UK Biobank recruited ≈500,000 people aged between 40-69 years in 2006-2010 from across the country to take part in this project ^83^. Potential participants were identified through National Health Service (NHS) registers according to being aged 40-69 and living within a reasonable travelling distance of an assessment centre. Assessment centres (22 in total) are located in accessible and convenient locations with a large surrounding population. Participants have undergone measures and provided samples and detailed information about themselves and agreed to have their health followed. Age was calculated from year and month of birth (day of month is missing, and was set to 1 for all subjects) to date of assessment.

##### Inclusion/Exclusion Criteria/Screening

Participants are excluded from scanning in the UKB according to fairly standard MRI safety/quality criteria, such as exclusions for metal implants, recent surgery, or health conditions directly problematic for MRI scanning, (e.g. problems hearing, breathing or extreme claustrophobia) ^84^. A generalized additive mixed model regressing area/thickness/volume nonlinear on age with random intercept per participant, was fitted independently at each of 163842 vertices. 40 observations were defined as outliers and excluded, as the absolute value of their residual was large than four times the residual standard error at more than 6000 vertices.

##### Variables Used for Birth weight and Education

Birth weight was extracted from UKB data field 20022. Units are in kilogram. If multiple birth weights were reported, we removed participants if the discrepancy between reported birth weights were greater than 10%, and we took the mean of the reported birth weights Education: For the Biobank participants’ generation, the UK school system provided free universal compulsory education between the ages of 5 and 15 to 16 years. Based on the UKB education data field 6138 (1 College or University degree; 2 A levels/AS levels or equivalent; 3 O levels/GCSEs or equivalent; 4 CSEs or equivalent; 5 NVQ or HND or HNC or equivalent; 6 Other professional qualifications eg: nursing, teaching; -7 None of the above; -3 Prefer not to answer, education was recoded to years using the following dictionary: edu_ukb_to_years = (1: 16, 2: 13, 3: 11, 4: 11, 5: 11, 6: 12, -7: 10, -3: np.NaN). If multiple education values were reported, the highest education value was chosen.

##### MRI scanning and processing

Imaging data were collected and processed by the <u>UK Biobank</u> (https://www.ukbiobank.ac.uk) as described in ^28^. Imaging data were collected using 3.0 T Siemens Skyra (32-channel head coil). Anatomical T1-weighted magnetization-prepared rapid gradient echo (MPRAGE) images were obtained in the sagittal plane at 1mm isotropic resolution, and T2 weighted FLAIR images were acquired at 1.05 x 1 x 1mm resolution in the sagittal plane. Images were processed by the UK Biobank using the FreeSurfer 6.0 software package.

## Specific information on image acquisition in the different samples

**Supplementary Table 1.**
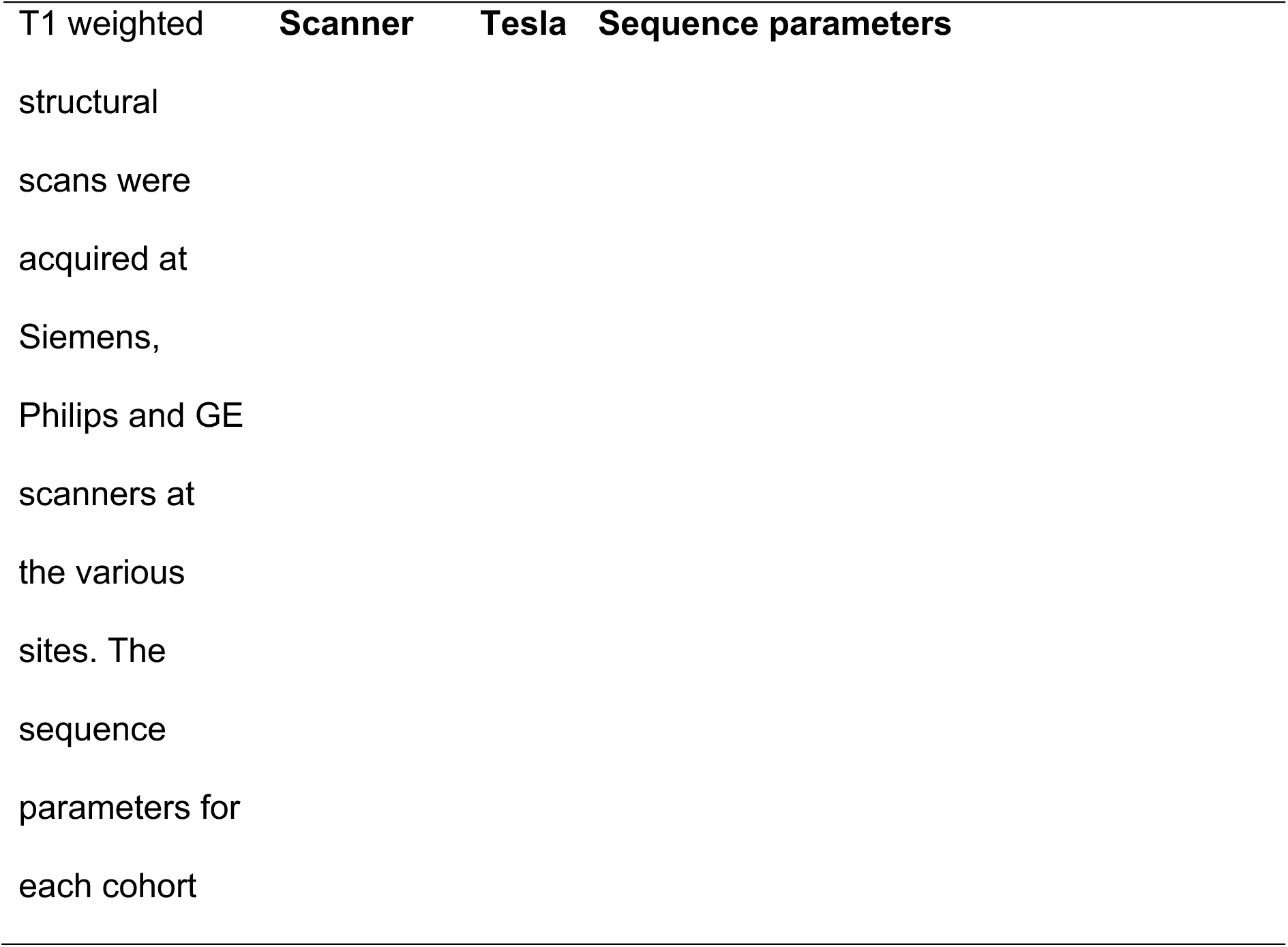
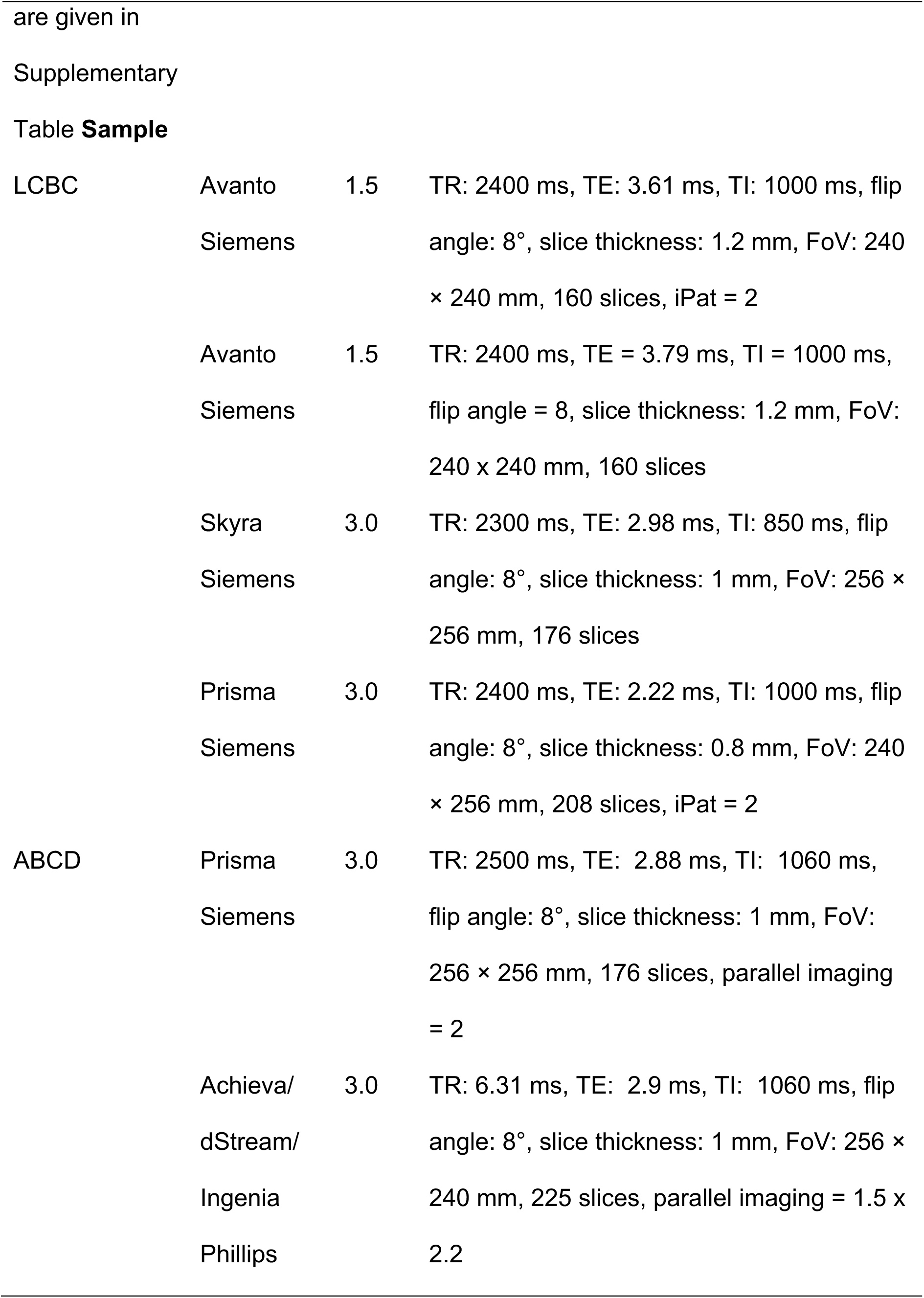
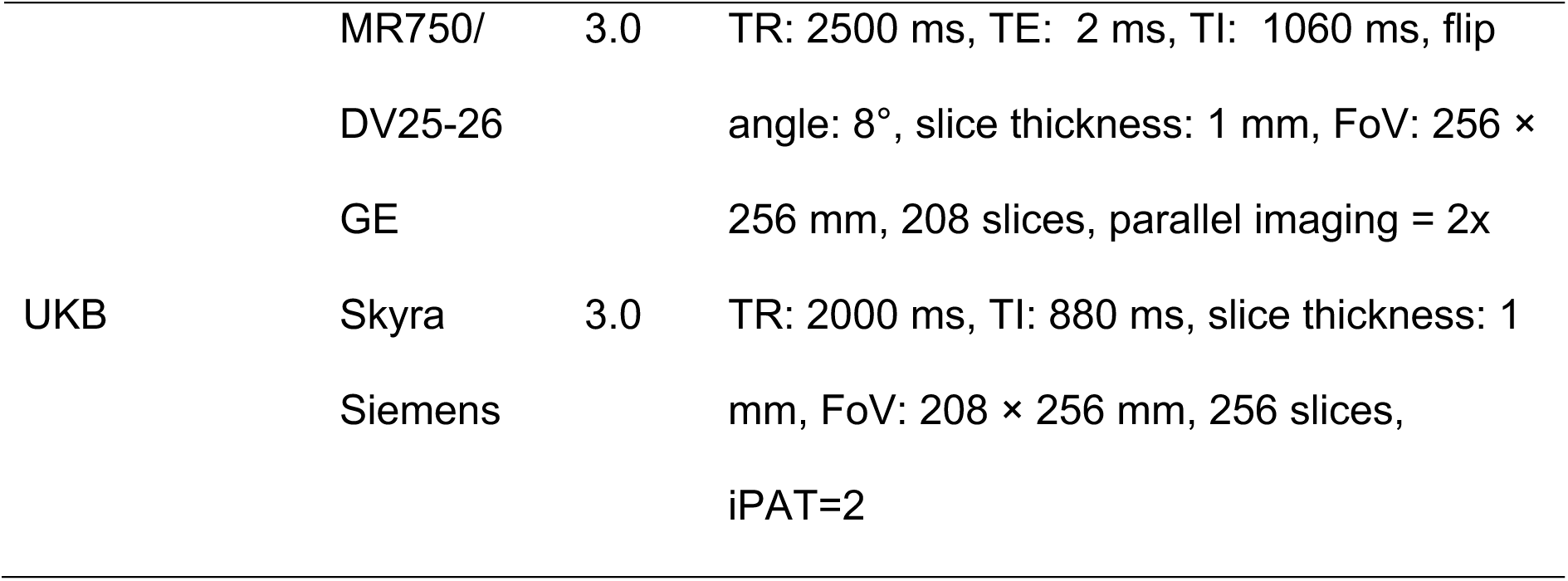
MR acquisition parameters. TR: Repetition time, TE: Echo time, TI: Inversion time, FoV: Field of View, iPat: in-plane acceleration, GRAPPA: GRAPPA acceleration factor. *Customized

## Supplementary Statistical methods

### Assessment of consistency of effects across and within samples

We assessed the spatial relationship between birth weight cortical correlates (βeta-maps) for cortical area, volume, and thickness across the different datasets (LCBC, UKB, ABCD) using Pearson’s correlations. Permutation-based significance testing (n = 10.000, p < 0.05, two-tailed, FDR-corrected) was performed using non-parametric spatial permutation models, i.e. spin tests as implemented by ^85,86^. Briefly, spin-tests generate spatially-constrained null distributions by applying random rotations to spherical projections of the brain. For each permutation, the original values at each coordinate are replaced with those of the closest rotated coordinate. Rotations are generated in one hemisphere and mirrored in the other.

Within-sample replicability was assessed in two different ways, an exploratory and a confirmatory analysis^37^. In the exploratory analysis, we assessed within-sample replicability by conducting the same vertex-wise cortical analysis in different subsamples. For each dataset and cortical measure, we assessed the effects of birth weight on cortical structure and cortical change |N| = 500 times using 50% of the original sample (participants’ split half). Beyond sample selection, all parameters remained identical as described in the *main* analysis (refer to Figure 1). Replicability – of a given vertex – was determined by the proportion of multiple comparison-corrected results across the different subsamples. In the confirmatory analysis, we assessed the proportion of significant vertices obtained in each *exploratory* (train) analysis that was also deemed significant – and in the same direction – in the remaining (test; 50%) subsample (p < 0.05 uncorrected). Subsamples without significant results were not considered (no results only in cortical thickness analyses). This is a criterion often followed for determining replicability^38^. Replicability analyses were performed on *fsaverage4* for computational reasons. Across samples replicability was performed as described in the within-sample replicability analysis (i.e., we assessed the exploratory and confirmatory replicability) except that split-half was not performed - the three datasets were compared with each other - and the analyses were performed in the original *fsaverage* space.

## Supplementary Results

**Supplementary Figure 1.**
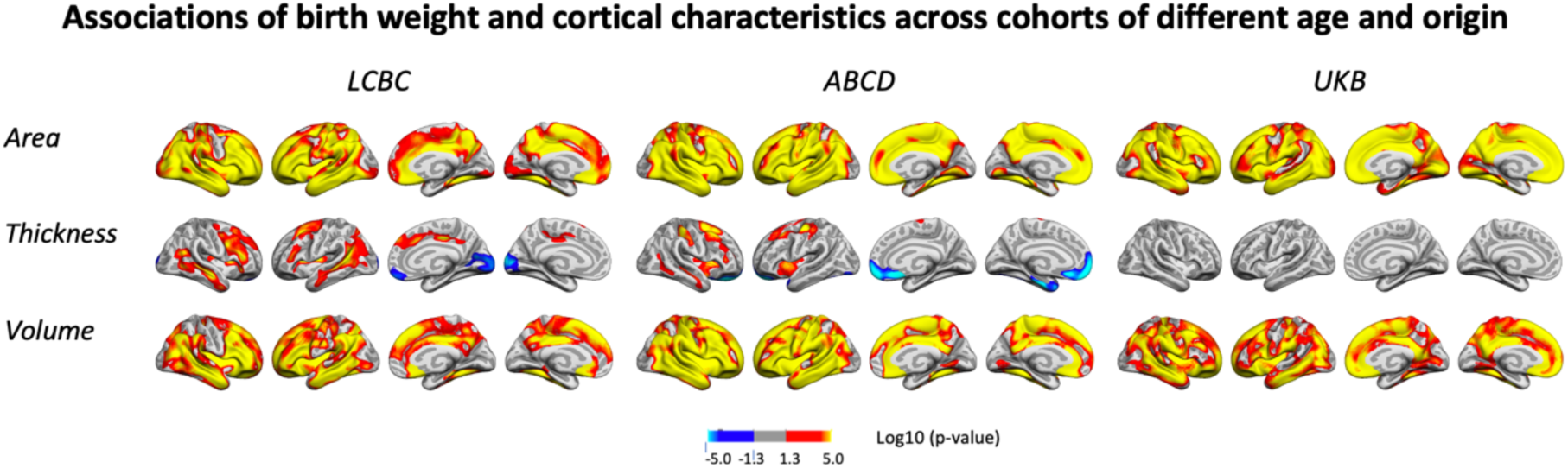
Relationships of birthweight and cortical area across LCBC, ABCD, and UKB samples when controlling for age, sex, time (interval since baseline) and scanner site (as well as ethnicity in the ABCD). Significant relationships are shown from left to right: lateral view, right and left hemisphere, and medial view, right and left hemisphere.

**Supplementary figure 2.**
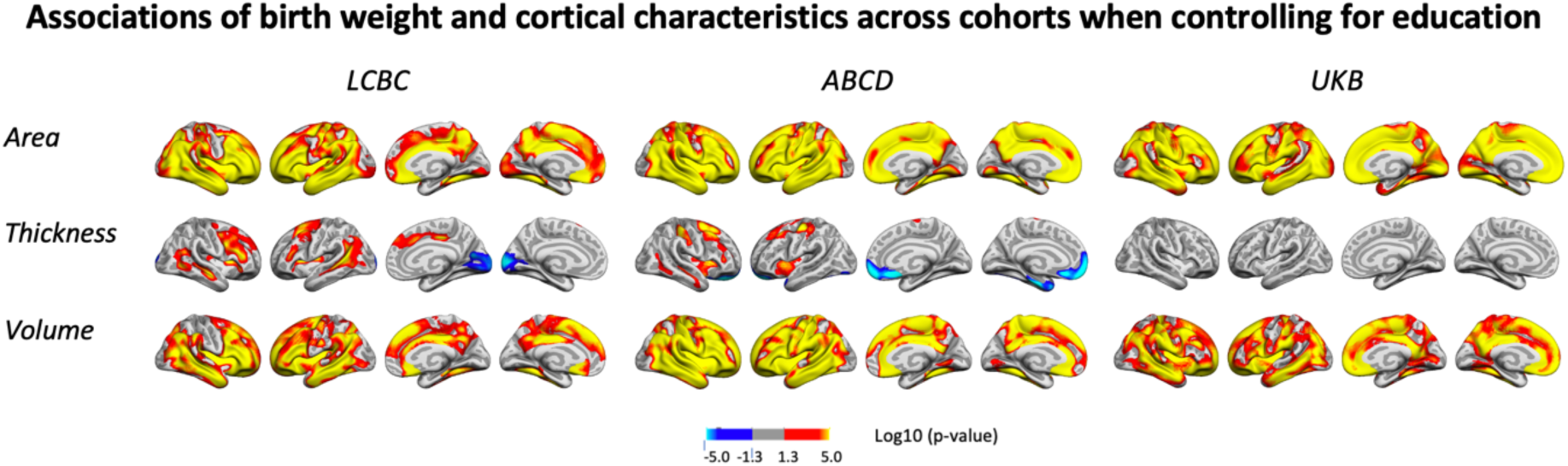
Relationships of birthweight and cortical characteristics across LCBC, ABCD, and UKB samples when controlling for education, age, sex, time (interval since baseline) and scanner site (as well as ethnicity in the ABCD). Significant relationships are shown from left to right: lateral view, right and left hemisphere, and medial view, right and left hemisphere.

**Supplementary figure 3.**
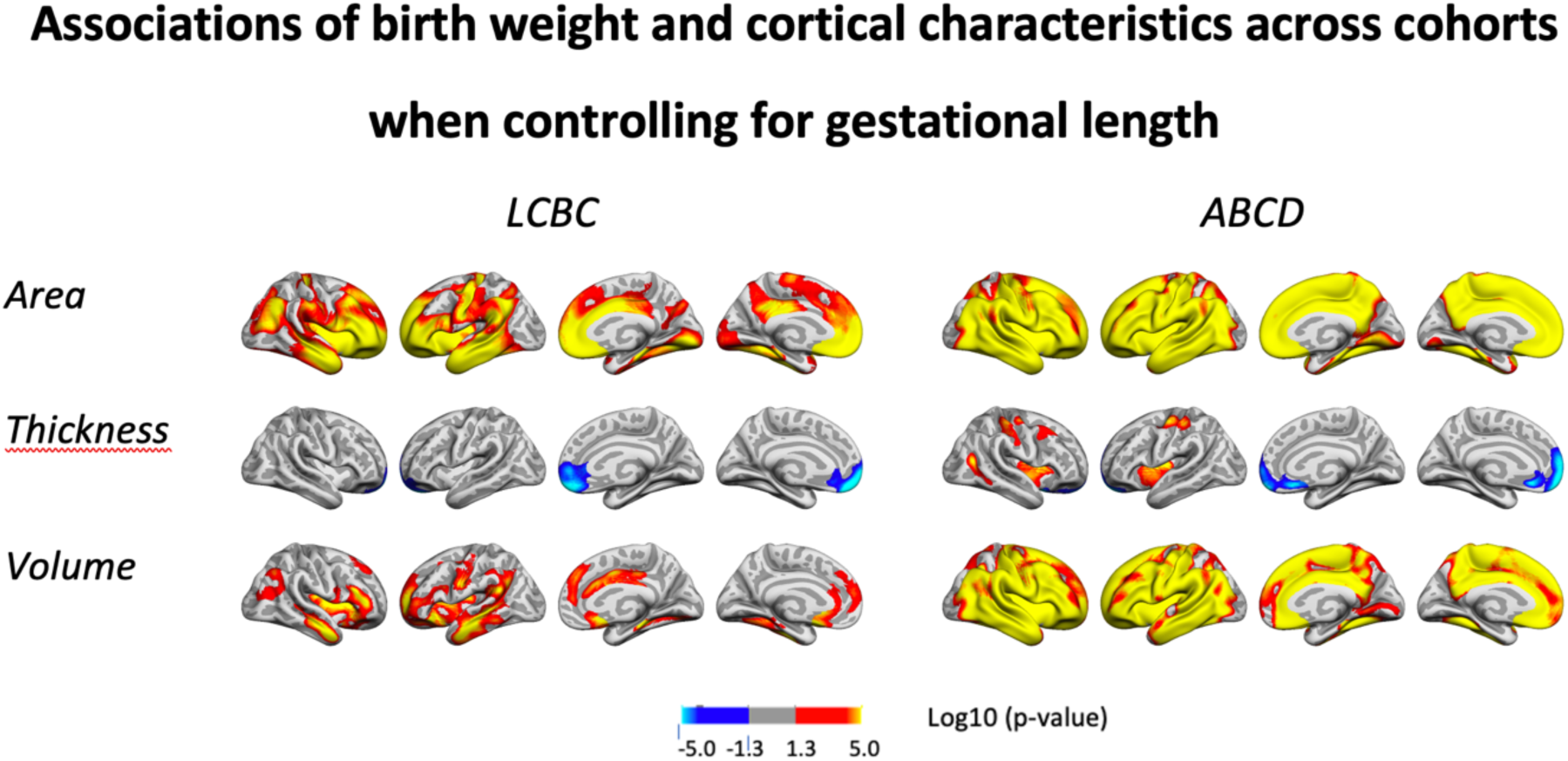
Relationships of birthweight and cortical characteristics across LCBC and ABCD samples when controlling for gestational length in weeks (LCBC) or weeks born prematurely (ABCD), age, sex, time (interval since baseline) and scanner site (as well as ethnicity in the ABCD). Significant relationships are shown from left to right: lateral view, right and left hemisphere, and medial view, right and left hemisphere.

**Supplementary Figure 4.**
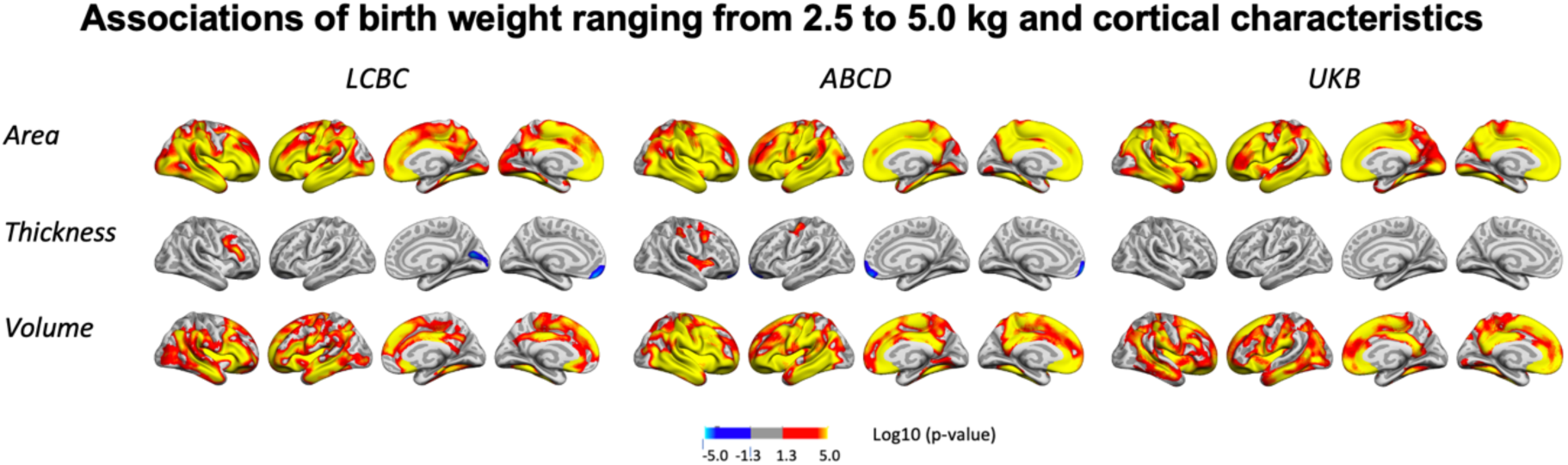
Relationships of birthweight and cortical characteristics across LCBC, ABCD, and UKB when restricting the samples to participants with birth weights between 2.5 and 5.0 kg, controlling for age, sex, time (interval since baseline) and scanner site (as well as ethnicity in the ABCD). (Compare to *Figure 1*.) Significant relationships are shown from left to right: lateral view, right and left hemisphere, and medial view, right and left hemisphere.

**Supplementary Figure 5.**
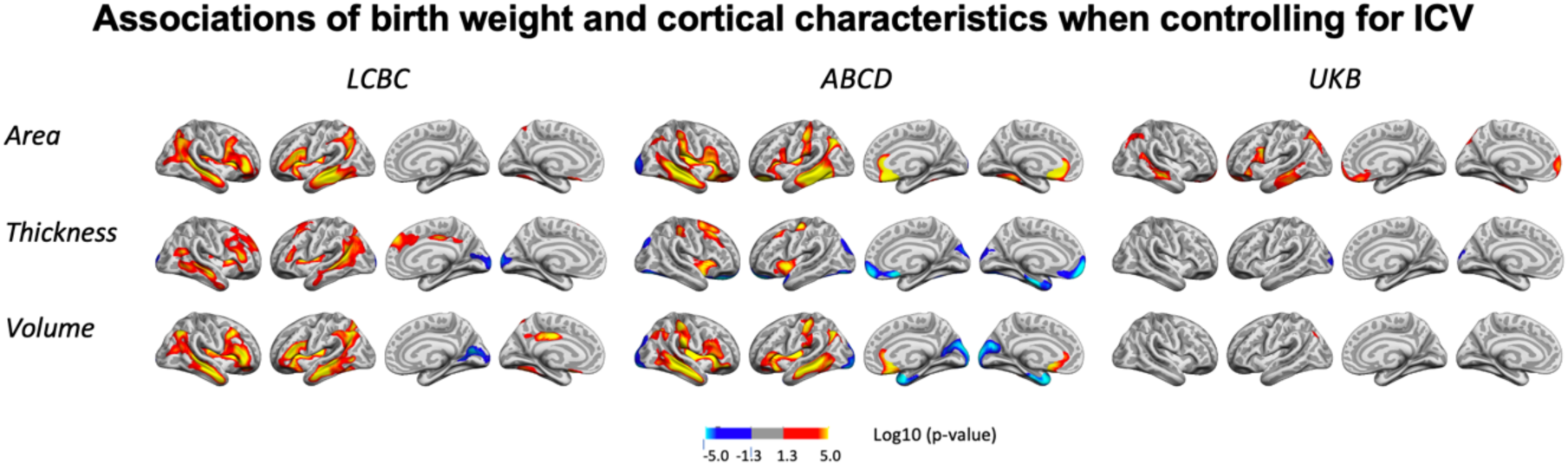
Relationships of birthweight and cortical characteristics across LCBC, ABCD, and UKB samples when controlling for intracranial volume (ICV), age, sex, time (interval since baseline) and scanner site (as well as ethnicity in the ABCD). Significant relationships are shown from left to right: lateral view, right and left hemisphere, and medial view, right and left hemisphere.

**Supplementary Figure 6.**
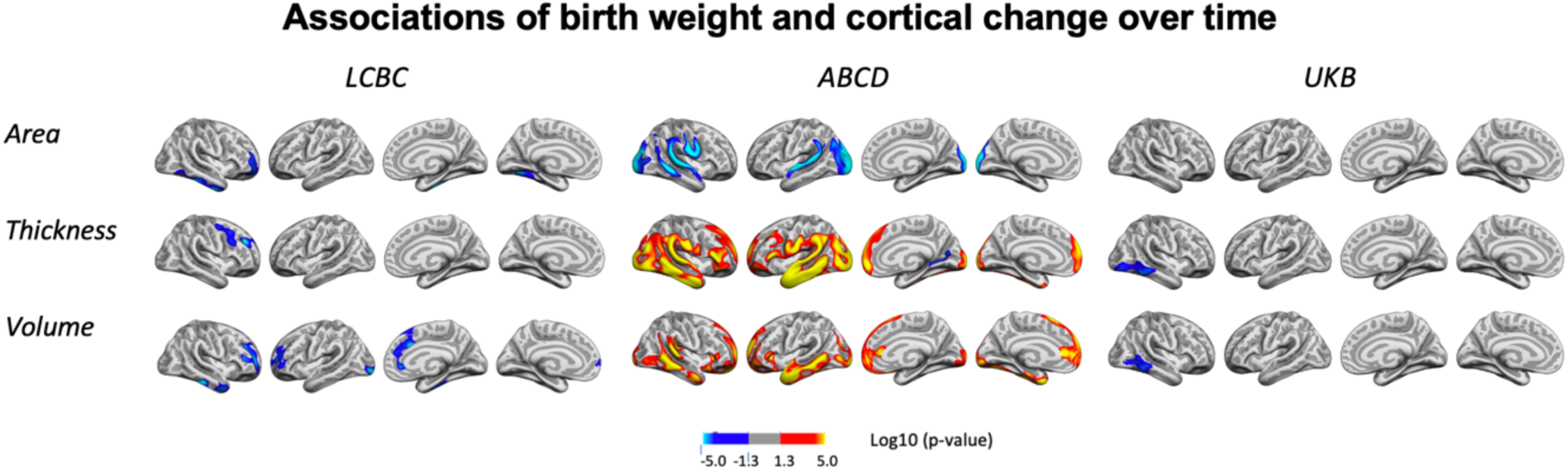
Interactions of BW and time on cortical characteristics across LCBC, ABCD, and UKB samples when controlling for age, sex, scanner site, time, birth weight, and the interaction of baseline age and time (as well as ethnicity in the ABCD). Significant relationships are shown from left to right: lateral view, right and left hemisphere, and medial view, right and left hemisphere.

**Supplementary Figure 7.**
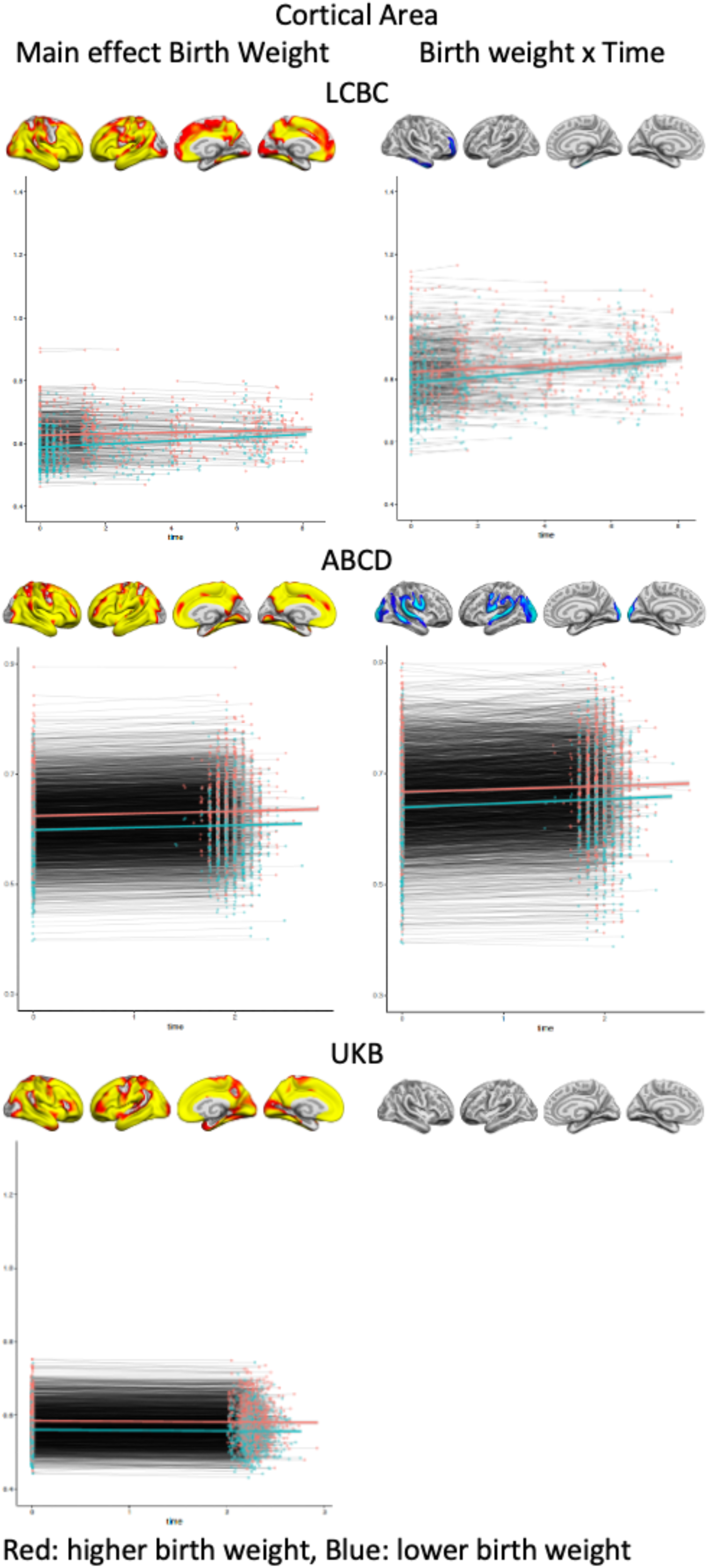
Plots showing individual data points and expected trajectories for cortical area within the significant regions (refer to *Figures 1 and 2*) of each sample split in two based on BW (higher BW in red color= upper half, lower BW in blue color= lower half of BW distribution) are shown. Y-axis: cortical area: mm2, thickness, X-axis: time in years.

**Supplementary Figure 8.**
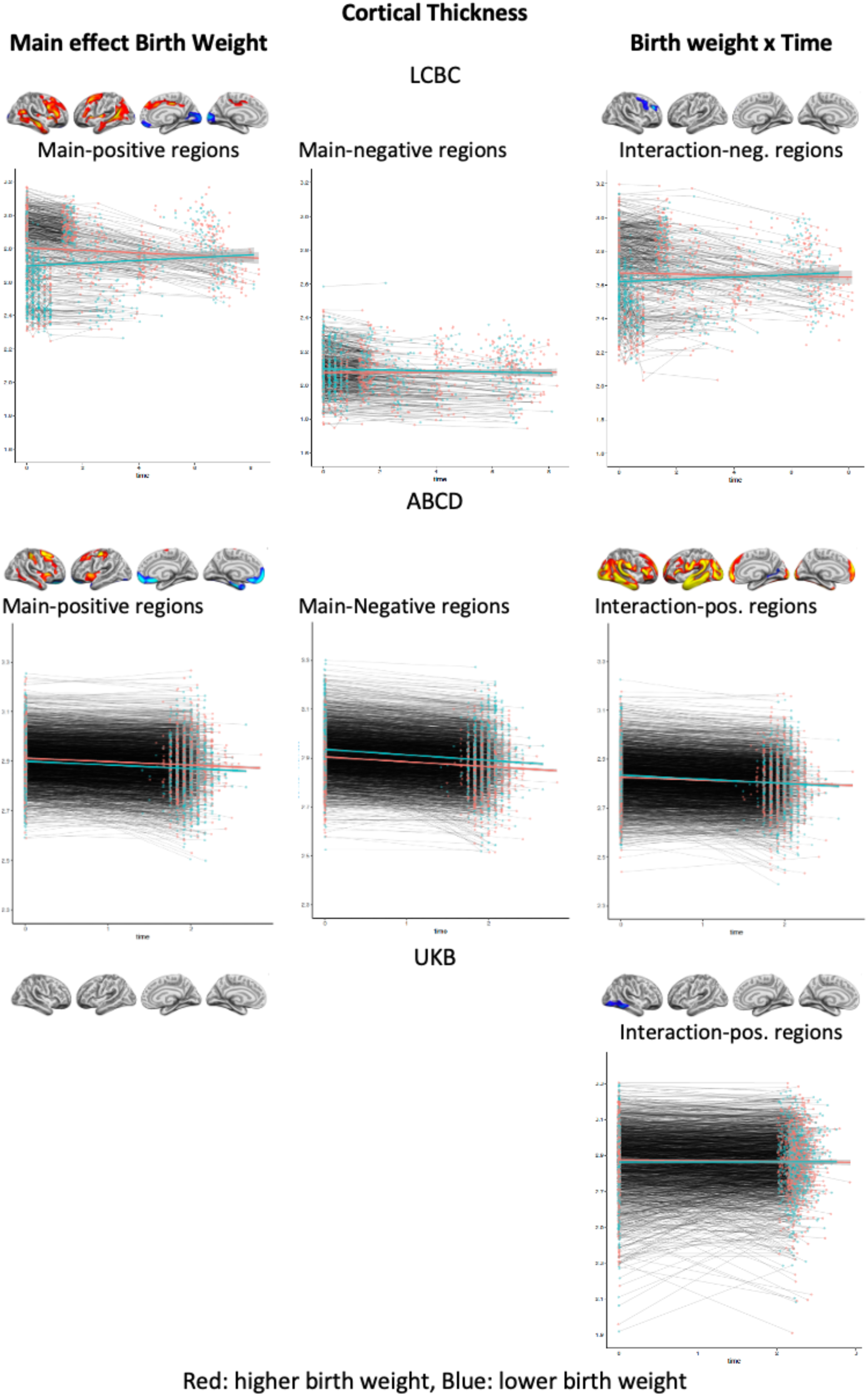
Plots showing individual data points and expected trajectories for cortical thickness within the significant regions (refer to *Figures 1 and 2*) of each sample split in two based on BW (higher BW in red color= upper half, lower BW in blue color= lower half of BW distribution) are shown. Y-axis: cortical thickness: mm, X-axis: time in years.

**Supplementary Figure 9.**
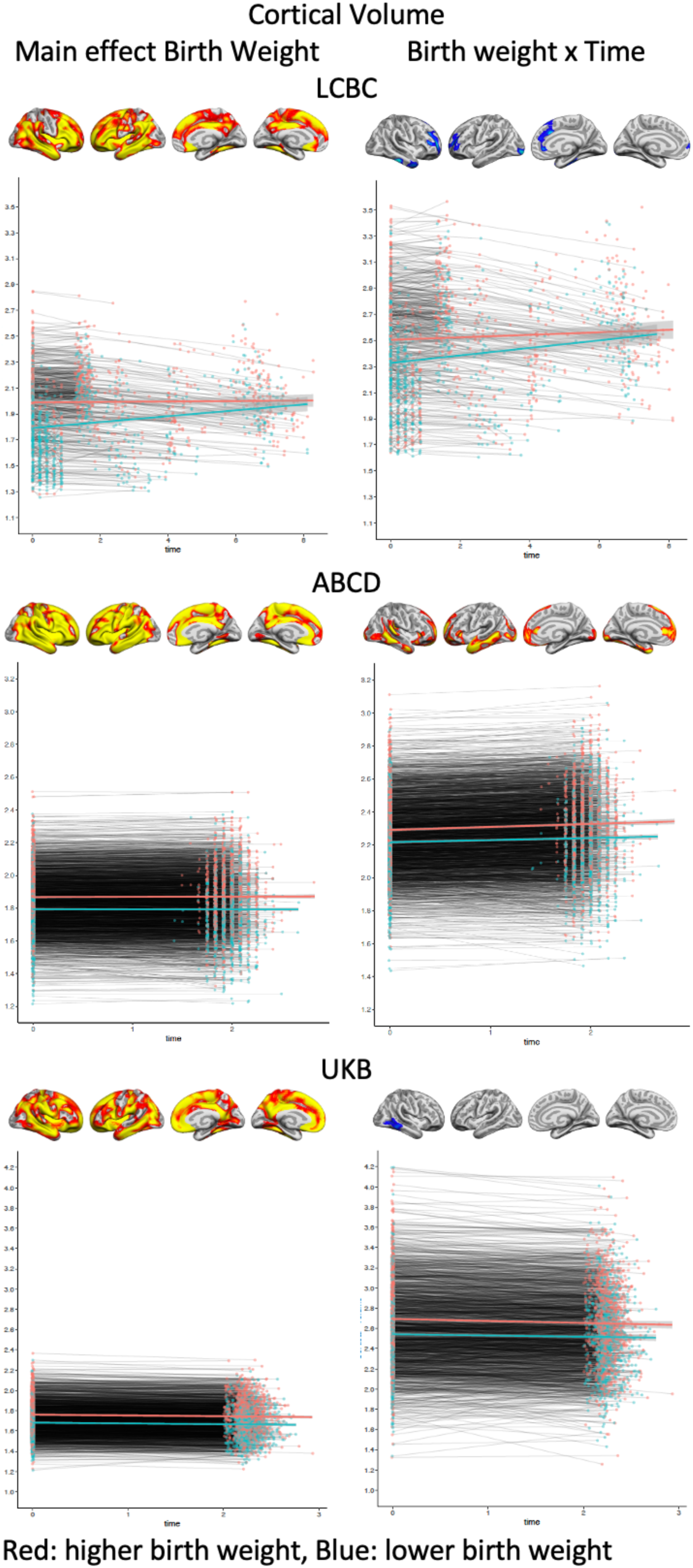
Plots showing individual data points and expected trajectories for cortical volume within the significant regions (refer to *Figures 1 and 2*) of each sample split in two based on BW (higher BW in red color= upper half, lower BW in blue color= lower half of BW distribution) are shown. Y-axis: cortical thickness: mm, X-axis: time in years.

**Supplementary Figure 10.**
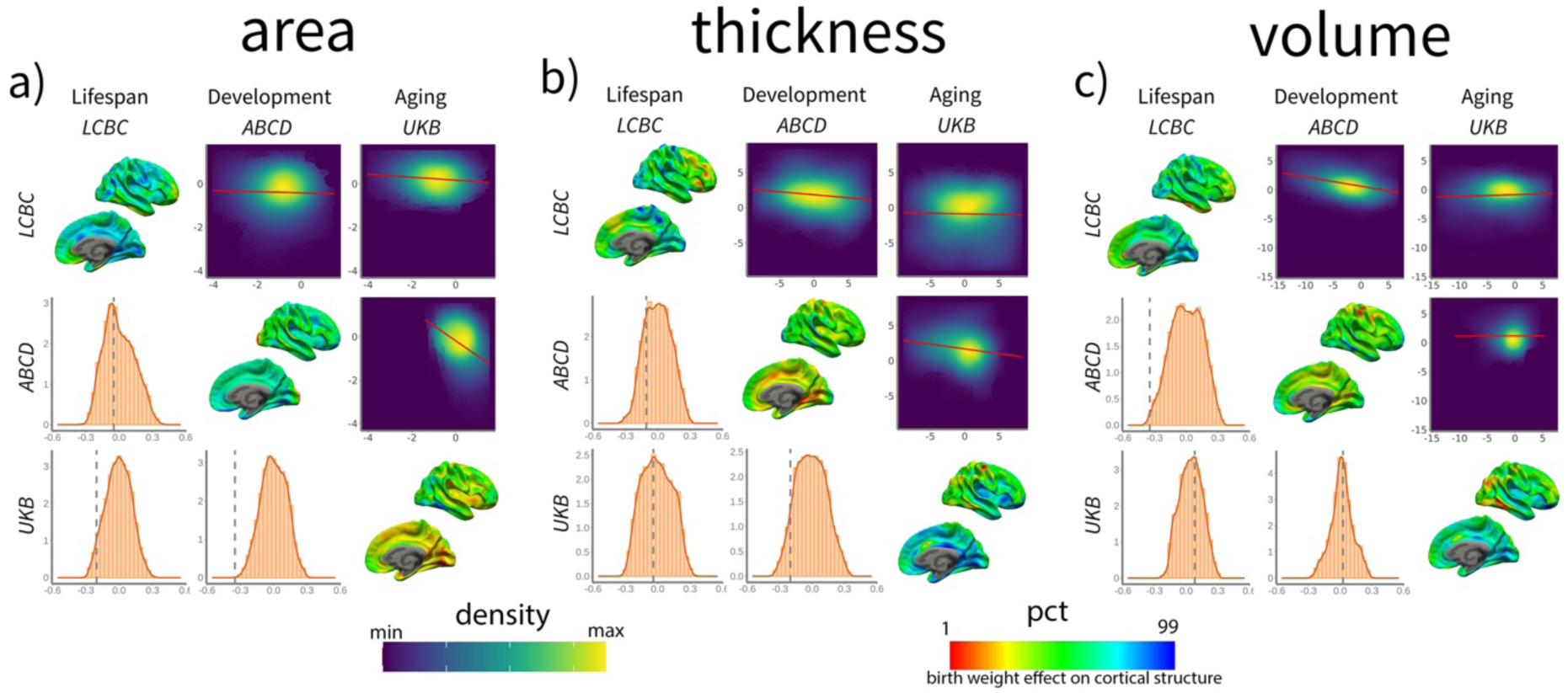
Spatial correlation of birth weight effects on brain structure change across datasets. For each panel, the upper triangular matrix shows Pearson’s (r) pairwise spatial correlation between the different cohorts’ cortical maps. Data is shown as a color-density plot. The red line represents the fitting between the two maps. The lower triangular matrix shows the significance testing. The dashed-grey line shows the empirical correlation, while the orange histogram represents the null distribution based on the spin test. The diagonal shows the effect of birth weight on cortical structure (right hemisphere shown only). Note that the βeta-maps are shown as a percentile red-green-blue scale, where red represents a lower (or more negative) effect of birth weight on cortical structure and vice versa. See Supplementary Table 2 for stats. The different panels show the spatial correlation of birth weight effects on cortical a) area, b) thickness, and c) volume. Units in the density maps represent birth weight effects as mm/g, mm^2^/g, and mm^3^/g (10e-5) for cortical thickness, area, and volume, respectively.

**Supplementary Table 2.**
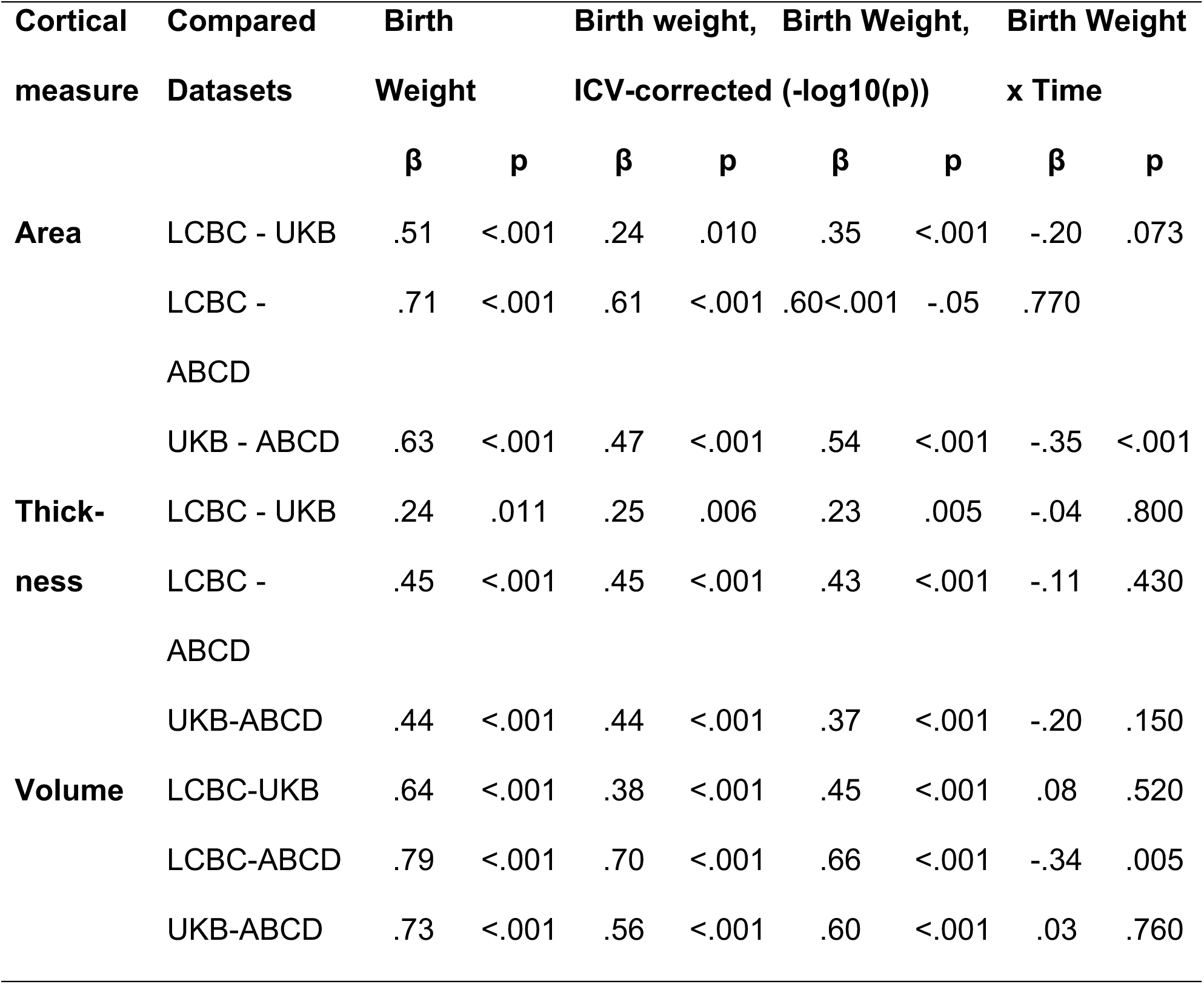
Spatial correlation of birth weight effects on brain structure across datasets. Pearson’s correlation and significance for pairwise spatial correlations between the different cohorts’ birth weight, with and without correction for ICV, and birth weight x time (years) -related cortical maps; i.e. effects of birth weight on cortical structure and change in cortical structure. Significance was assessed using the spin test. Spatial correlations were assessed using the Beta maps (and -log10(p) for birth weight). See Figure 3 and Supplementary Figure 9 and 10 for a visual representation. P-values are FDR-corrected (n = 9).

The degree of within-sample replicability was high for both area and volume but not for cortical thickness (see **Supplementary Figure 11**) in the three datasets (LCBC, ABCD, and UKB). For area and volume, some vertices were fully replicable, i.e., significant results were found in this vertex regardless of sample variability. *Exploratory* replicability [median (interquartile range); LCBC, ABCD, UKB] was .46 (.56), .85 (.47), .89 (.43) for area, .36 (.47), .50 (.65), .79 (.68) for volume, and .008 (.04), .00 (.00), .002 (.02) for thickness. *Exploratory* replicability indexes for each voxel the proportion of significant results found using different |N| = 500 subsamples. The confirmatory replicability analyses also showed good replicability for area and volume but not for cortical thickness. Confirmatory replicability indexes the proportion of voxels (per analysis) that was significant both in the exploratory subsample (p < 0.01, FWE-corrected) and in the test subsample (p < 0.05). The proportion of vertices that were significant both in the *exploratory* and *test* subsamples was .74 (.17), .91 (.08), .93 (.08) for area, .68 (.19), .84 (.12), .92 (.09) for volume, and .28 (.28), .00 (.00), .58 (.18) for thickness.

The *exploratory* replicability of birth weight on cortical change was negligible across datasets and measures [.00 (.00), .00 (.00), .00 (.00) for area, .02 (.09), .00 (.02), .01 (.03) for volume, and .01 (.05), .01 (.14), .00 (.01) for thickness] while confirmatory replicability was generally poor, with the exception of the ABCD dataset [.02 (.05), .68 (.35), .00 (.00) for area, .08 (.14), .56 (.25), .00 (.02) for volume, and .37 (.26), .60 (.27), .01 (.03) for thickness] (see **Supplementary Table 3**).

These results are not fully comparable to other studies assessing the replicability of brain-phenotype associations due to analytical differences (e.g. sample size, multiple-comparison correction method)^20,37^, yet clearly show that the rate of replicability of BW associations with cortical area and volume are comparable to benchmark brain-phenotype associations such as body-mass index and age^69^. Lower levels of replicability in the LCBC subsample are likely attributable to higher sample variability (e.g. increased age span). Kinship may lead to inflated patterns of replicability within the ABCD cohort. Confirmatory replicability is, also, to some degree, affected by sample size, and thus the estimates of confirmatory replicability may be somewhat inflated in the ABCD dataset.

Finally, the degree of across-sample replicability was high for the effects of birth weight on cortical area and volume (average confirmatory replicability = .96 and .93), low for thickness (.27), and negligible for the effects of birth weight on cortical change (.03, .06, and .06). See further information in **Supplementary Table 4**.

**Supplementary Figure 11.**
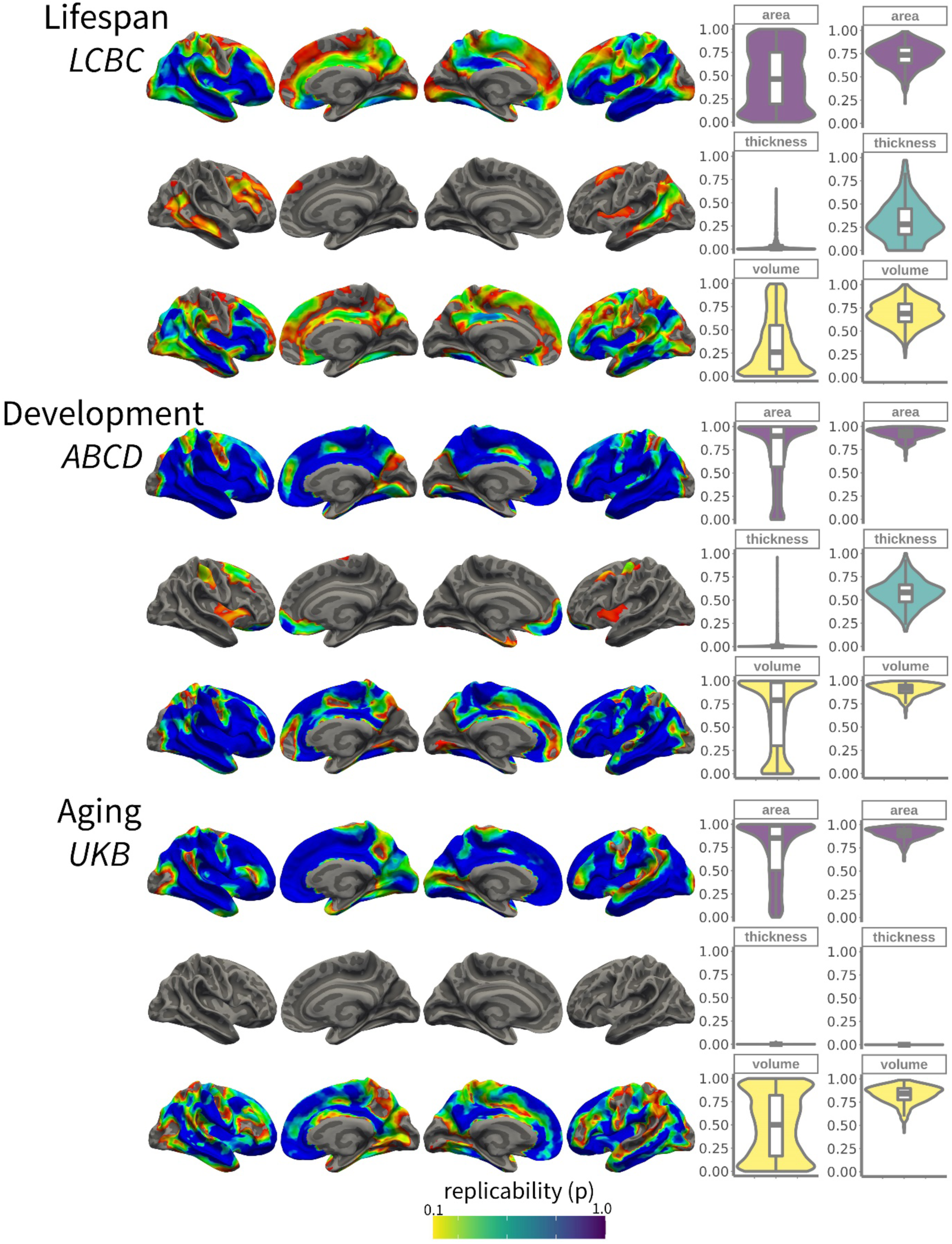
The degree of within-sample replicability of birth weight effects on cortical structure for LCBC, ABCD, and UKB. Brain images show the exploratory replicability analyses; i.e. the proportion of results (refer to Figure 1) that are significant across different subsamples (50% of the original sample; |N| = 500 subsamples). The overlay is thresholded at replicability (p) = .1. For visual purposes data has been up-sampled and smoothed (FWHM = 5) on fsaverage space. Left-side violin plots show the exploratory replicability; i.e., the proportion of significant results across the different subsamples (unit = voxel). Right-side plots show the confirmatory replicability; i.e. for each sample, the proportion of results (p < 0.01; FWE-corrected) that also pass significance criteria (p < 0.05) in a test subsample (|N| = 500 subsamples) (unit = analysis). Embedded boxplots display median, interquartile range, and range.

**Supplementary Figure 12.**
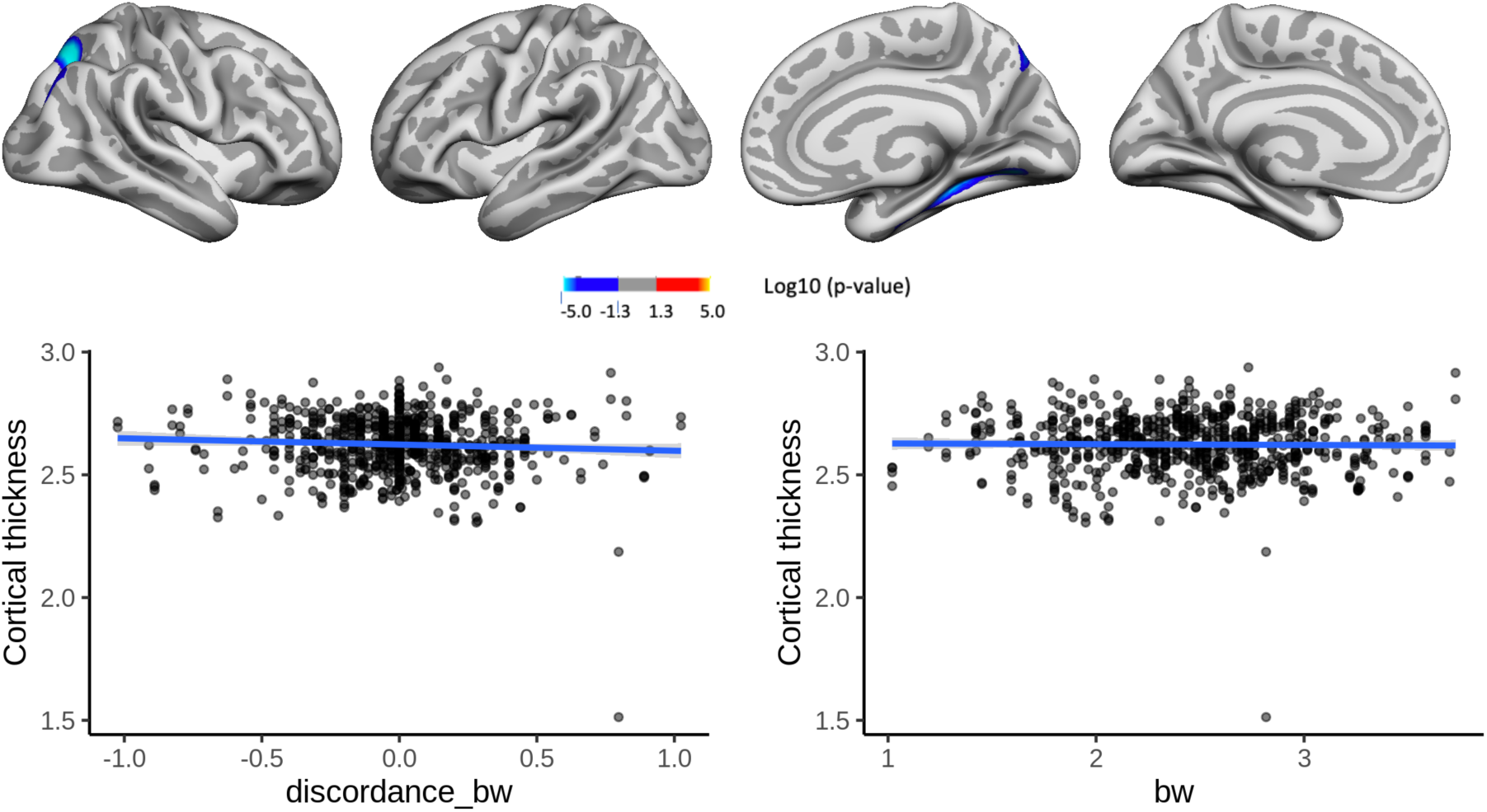
Interaction effects of birth weight discordance and time on cortical thickness in the sample of monozygotic (MZ) twins. Significant relationships are shown from left to right: lateral view, right and left hemisphere, and medial view, right and left hemisphere. Plots showing individual data points and expected trajectories for cortical thickness in mm (Y-axes) within the significant regions according to BW discordance (left panel) and BW (right panel) in kilograms (X-axes) are shown.

**Supplementary Figure 13.**
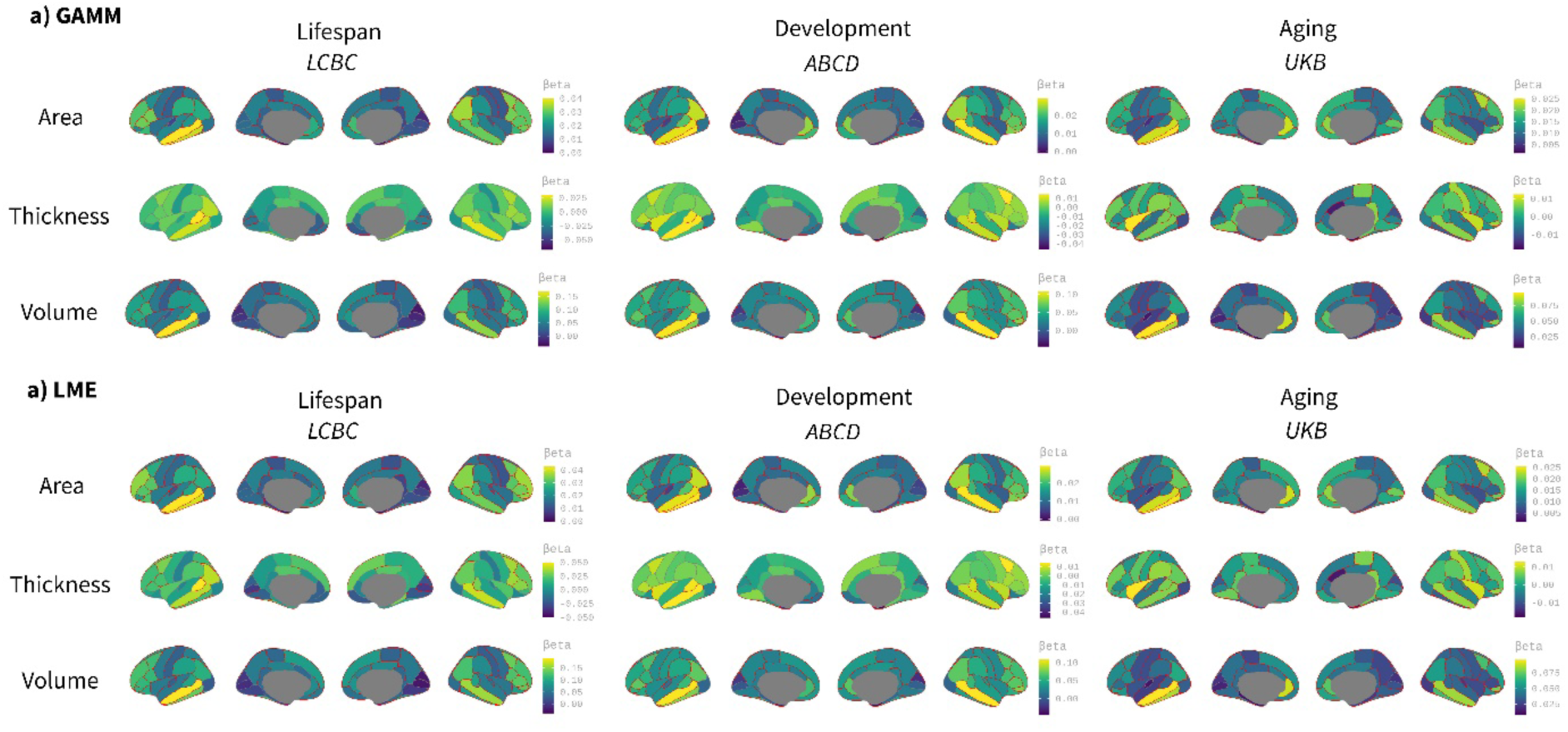
Comparison between spline (GAMM) and linear (LME) models on the effect of birth weight on cortical characteristics. Age was fitted either as a smoothing spline using generalized additive mixed models (GAMM, mgcv r-package) or a linear regressor with a linear mixed models (LME, lmer r-package) framework. The analyses were performed on ROI-based using the Desikan-Killiany atlas. Significance was considered at a FDR corrected threshold of p < 0.04. All the remaining parameters were comparable to the main analyses shown in Figure 1. The viridis-yellow scale represents the lower-higher Beta regressors. Red contour displays regions showing significant effects of birth weight. Note the high correspondence with both fitting models. Differences are only noticeable in the LCBC sample due to the datasets’ wider age range (i.e., lifespan dataset).

**Supplementary Figure 14.**
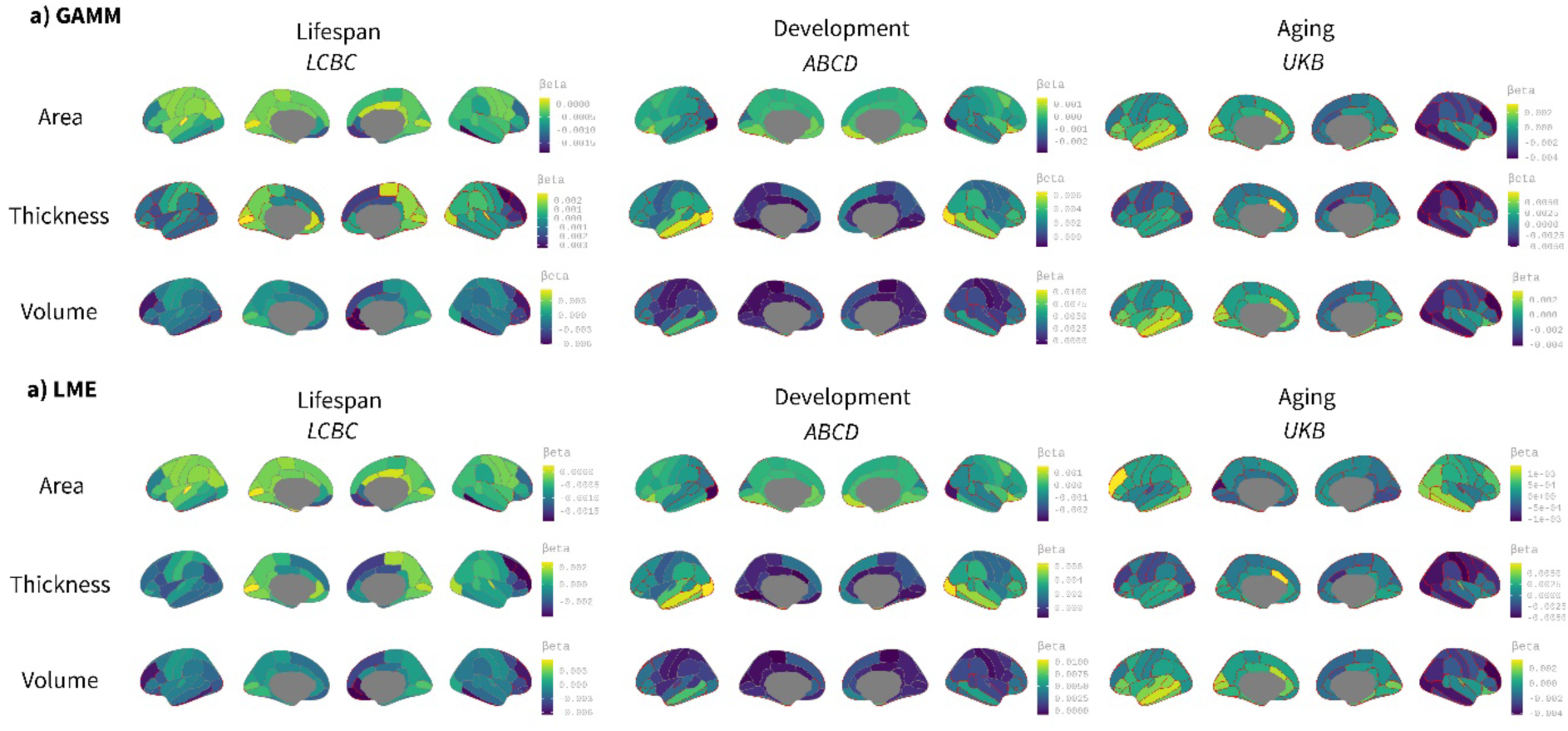
Comparison between spline (GAMM) and linear (LME) models on the effect of birth weight on cortical change. Age was fitted either as a smoothing spline using generalized additive mixed models (GAMM, mgcv r-package) or a linear regressor with a linear mixed models (LME, lmer r-package) framework. The analyses were performed on ROI-based using the Desikan-Killiany atlas. Significance was considered at a FDR corrected threshold of p < 0.04. All the remaining parameters were comparable to the main analyses shown in Figure 1. The viridis-yellow scale represents the lower-higher Beta regressors. Red contour displays regions showing significant effects of birth weight. Note the high correspondence with both fitting models. Differences are only noticeable in the LCBC sample due to the datasets’ wider age range (i.e., lifespan dataset).

**Supplementary Table 3.**
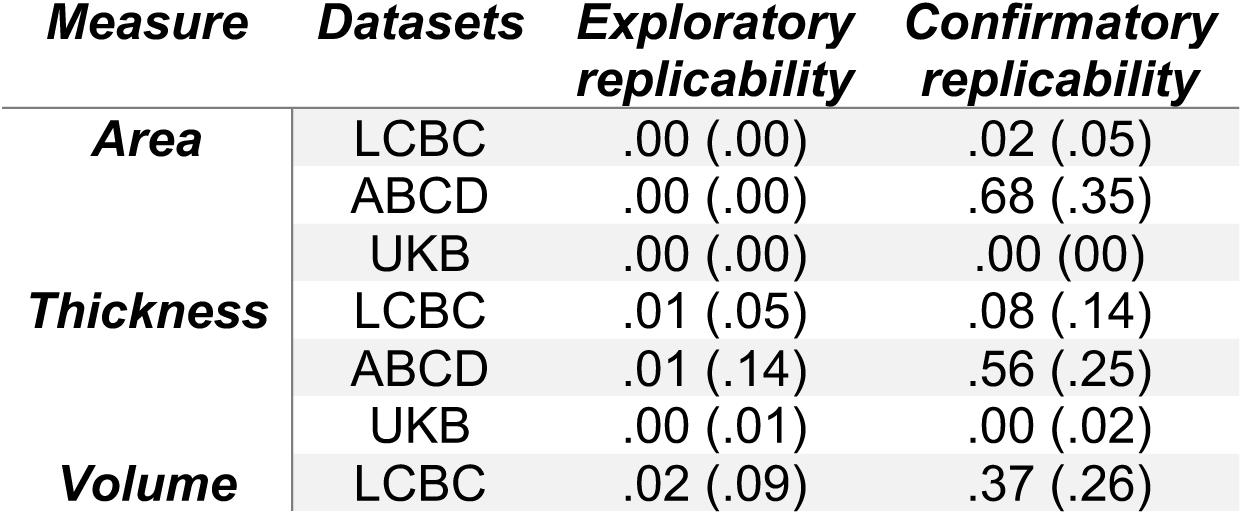
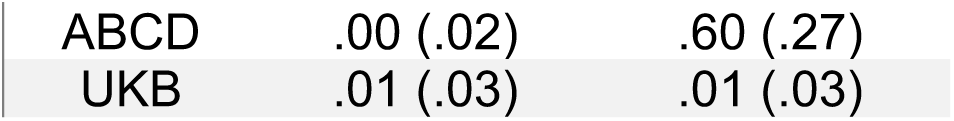
Exploratory and confirmatory replicability of birth weight on cortical change within datasets. Units represent median and interquartile range.

**Supplementary Table 4.**
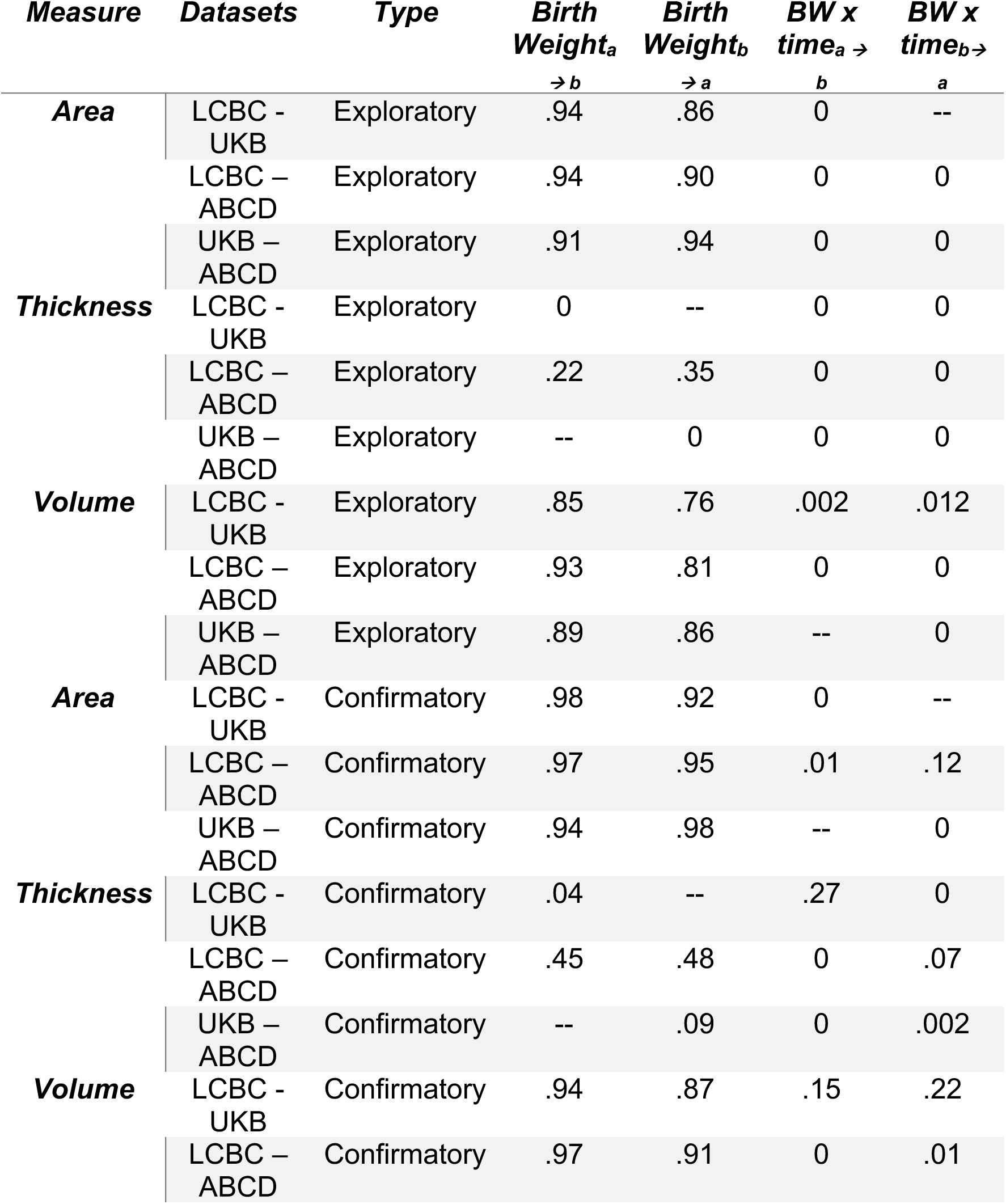
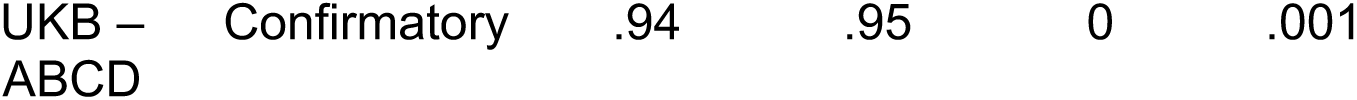
Exploratory and confirmatory replicability across datasets. “--" denotes no significant clusters in the right-hand dataset. ^a → b^ and _b → a_ denotes directionality of the replicability analyses being “a” and “b” in the left and right hand of the “Compared Datasets” column. BW = Birth weight.

| *Measure* | *Datasets* | *Type* | *Birth Weight<sub>a</sub>*<br>→ *b* | *Birth Weight<sub>b</sub>*<br>→ *a* | *BW x time<sub>a</sub>* →<br>*b* | *BW x time<sub>b</sub>* →<br>*a* |
| --- | --- | --- | --- | --- | --- | --- |
| **Area** | LCBC - UKB | Exploratory | .94 | .86 | 0 | -- |
|  | LCBC – ABCD | Exploratory | .94 | .90 | 0 | 0 |
|  | UKB – ABCD | Exploratory | .91 | .94 | 0 | 0 |
| **Thickness** | LCBC - UKB | Exploratory | 0 | -- | 0 | 0 |
|  | LCBC – ABCD | Exploratory | .22 | .35 | 0 | 0 |
|  | UKB – ABCD | Exploratory | -- | 0 | 0 | 0 |
| **Volume** | LCBC - UKB | Exploratory | .85 | .76 | .002 | .012 |
|  | LCBC – ABCD | Exploratory | .93 | .81 | 0 | 0 |
|  | UKB – ABCD | Exploratory | .89 | .86 | -- | 0 |
| **Area** | LCBC - UKB | Confirmatory | .98 | .92 | 0 | -- |
|  | LCBC – ABCD | Confirmatory | .97 | .95 | .01 | .12 |
|  | UKB – ABCD | Confirmatory | .94 | .98 | -- | 0 |
| **Thickness** | LCBC - UKB | Confirmatory | .04 | -- | .27 | 0 |
|  | LCBC – ABCD | Confirmatory | .45 | .48 | 0 | .07 |
|  | UKB – ABCD | Confirmatory | -- | .09 | 0 | .002 |
| **Volume** | LCBC - UKB | Confirmatory | .94 | .87 | .15 | .22 |
|  | LCBC – ABCD | Confirmatory | .97 | .91 | 0 | .01 |

**Supplementary Table 4.** Exploratory and confirmatory replicability across datasets.
|  |  |  |  |  |  |
| --- | --- | --- | --- | --- | --- |
| UKB – | Confirmatory | .94 | .95 | 0 | .001 |
| ABCD |  |  |  |  |  |

